# Compliant DNA Origami Nanoactuators as Size-Selective Nanopores

**DOI:** 10.1101/2024.04.12.589171

**Authors:** Ze Yu, Anna V. Baptist, Susanne C. M. Reinhardt, Eva Bertosin, Cees Dekker, Ralf Jungmann, Amelie Heuer-Jungemann, Sabina Caneva

**Affiliations:** Department of Precision and Microsystems Engineering, Delft University of Technology, Mekelweg 2, 2628 CD, Delft, The Netherlands; Max Planck Institute of Biochemistry, Am Klopferspitz 18, 82152 Martinsried, Bavaria, Germany and Center for NanoScience, Ludwig-Maximilians-Universität, Geschwister-Scholl-Platz 1, 80539 Munich, Bavaria, Germany; Faculty of Physics, Ludwig-Maximilians-Universität, Geschwister-Scholl-Platz 1, 80539 Munich, Bavaria, Germany; Department of Bionanoscience, Kavli Institute of Nanoscience, Delft University of Technology, 2629 HZ Delft, The Netherlands

**Keywords:** DNA origami, nanopores, compliant mechanism, DNA PAINT, nanoactuator

## Abstract

Biological nanopores crucially control the import and export of biomolecules across lipid membranes in cells. They have found widespread use in biophysics and biotechnology, where their typically narrow, fixed diameters enable selective transport of ions and small molecules as well as DNA and peptides for sequencing applications. Yet, due to their small channel sizes, they preclude the passage of large macromolecules, e.g., therapeutics. Here, we harness the unique combined properties of DNA origami nanotechnology, machine-inspired design, and synthetic biology, to present a structurally reconfigurable DNA origami MechanoPore (MP) that features a lumen that is tuneable in size through molecular triggers. Controllable switching of MPs between three stable states is confirmed by 3D-DNA-PAINT super-resolution imaging and through dye-influx assays, after reconstitution of the large MPs in the membrane of liposomes via an inverted-emulsion cDICE technique. Confocal imaging of transmembrane transport shows size-selective behaviour with adjustable thresholds. Importantly, the conformational changes are fully reversible, attesting to the robust mechanical switching that overcomes pressure from the surrounding lipid molecules. These MPs advance nanopore technology, offering functional nanostructures that can be tuned on-demand – thereby impacting fields as diverse as drug-delivery, biomolecule sorting and sensing, as well as bottom-up synthetic biology.

## Introduction

Transmembrane nanopores are nanoscale channels that span biological membranes and play a vital role in many biological processes, such as cell communication and signal transduction.[1] Built from pore-forming proteins, biological nanopores have been repurposed with great success for biotechnological applications, such as DNA and protein sequencing,[2] sensing of biomarkers and post-translational modifications (PTMs),[3] monitoring of reaction trajectories, and small molecule delivery.[4] However, their small (typically <5 nm) and fixed diameters restrict their versatility and therefore application range. The size limitations preclude sensing and transmembrane transport of large macromolecules, for example clinically relevant polymers and proteins. Thus, the generation of nanopores with wider, tuneable diameters will expand their application profile and help address challenges in macromolecule delivery, as well as provide a platform for protein (dis-)aggregation studies and synthetic cell research. For example, by mimicking components such as the nuclear pore complex, which in nature consist of large (>40 nm) channel diameters,[5] they would allow better understanding and bottom-up engineering of the selective transport of cargo across the nuclear envelope.

A breakthrough in this direction has been the emergence of synthetic nanopores, which are expanding the field of nanopore technology by providing a pathway to overcome the size restrictions of naturally occurring protein pores.[1, 6] De novo design offers an opportunity to generate nanopores with tailored size, shape, and functionalities far beyond their biological counterparts. [7–12]

In recent years, DNA has become a very attractive building material for the bottom-up design of diverse nanoarchitectures, including nanopores.[8] In particular, the DNA origami technique allows for the precise self-assembly of nanostructures by folding a long single-stranded DNA (ssDNA) scaffold with many short DNA oligonucleotides (‘staples’), into designer 2D and 3D shapes.[13–14] The programmability and molecular resolution of this approach enables the formation of transmembrane nanopores with precise geometries and sizes, as well as controllable gating mechanisms. DNA origami-based nanopores with lumens ranging from 5 nm to 35 nm in diameter have been demonstrated in a number of previous works.[8–9, 15–23] Additional functionalities, such as gating via DNA locking/unlocking strands,[24] opening/closing a ‘lid’,[12] or reversibly attaching polymer brushes inside the channel to block biomolecule translocation,[22] have also been reported. However, to date, most reported DNA origami nanopores have been static, meaning that while access to the channel can be controlled, the diameter cannot be modulated on-demand. It remains challenging to realize fully reconfigurable nanopores capable of controlled and reversible conformational changes in both expansion and contraction (opening and closing), and with fully retained functionality after incorporation into operationally relevant environments, e.g. in a lipid membrane. A first step towards this was recently demonstrated in a responsive, semi-flexible DNA origami nanopore that adapted to biochemical (DNA and protein binding) and physical (transmembrane voltage) stimuli. However, this mainly resulted in irreversible opening or closing of the nanopores, somewhat limiting their applicability.[25] Thus, the formation of a fully reconfigurable DNA nanopore with a large lumen size and controlled, reversible initiation and stabilization of different, pre-designed states, able to function in liposomes, yet remains to be realized.

A promising route towards the fabrication of such fully controllable artificial transmembrane channels from DNA origami is the implementation of a compliant mechanism for the pre-programmed opening and closing of the nanopores. Such compliant mechanisms are used to achieve force and motion via the elastic deformation of the components of a flexible device.[26–30] They have been realized experimentally in DNA origami nanoactuators, showing that nanodevices made from DNA can undergo large conformational changes in a precise and controlled manner.[31]

Here, we introduce such a compliant mechanism to design a fully reconfigurable transmembrane DNA origami nanopore, hereafter referred to as the MechanoPore (MP). This MP is capable of controllably switching between three configurations (closed ↔ intermediate ↔ open) in response to an external trigger, with resulting nanopore areas ranging from ∼100 nm^2^ to 500 nm^2^. The incorporation of flexible ssDNA linkers and struts in the design allows for the fully reversible switching of the nanopore (via strand addition and displacement) as well as the controlled stabilization of defined open and closed states, resulting in a high level of control. Importantly, the switching mechanism implemented here enables the controlled operation of the nanopores even after their successful insertion into the lipid membranes of giant unilamellar vesicles (GUVs), enabling on-demand translocation of molecules through the pore that is size-selective, i.e. allowing small molecules to pass while blocking transport of large molecules. Our results clearly show that the very same DNA origami nanopores can allow or block the transmembrane passage of macromolecules of certain sizes depending on their adjusted configuration, demonstrating their potential for precisely controlled size-selective molecular transport across lipid membranes. This work paves the way towards the development of DNA-based nanopores for biomedical applications, including targeted drug delivery and molecular sorting, as well as to gateable communication channels in synthetic cell research.

## Results and Discussion

### Design and assembly of a compliant DNA origami nanoactuator

Our mechanically actuatable MP consists of a rhombic structure composed of 4 flipped ‘L’-shaped subunits made up of 22 helix bundles designed in a honeycomb lattice in order to minimize global twist (see **Figures S1 & S2** for caDNAno[32] layout). In a subunit, the bottom part forms the transmembrane section that embeds into the lipid bilayer (barrel), and the upper part rests on the top of the membrane (cap). Switchable conformations are realized by the incorporation of flexible ssDNA segments made from scaffold parts connecting the adjacent rigid subunits, resulting in some flexibility in the corner regions but limiting out-of-plane movement (**Figure 1a-c**). This design feature allows for full control over the nanopore’s configuration via the addition of trigger strands, enabling an active opening and closing mechanism, where the final states are fully stabilized (**Figure 1d**). In the absence of a trigger strand, the MP is in an ‘intermediate’ state with an opening angle of around 50° and a diameter of the enclosed lumen of around 20 nm. The addition of trigger strands (‘opening strands’) complementary to the ssDNA regions inside the MP results in the formation of double-stranded struts with increased stiffness, causing a conformational change into the open, square-shaped state with an opening angle of around 90° and a significantly increased inner lumen (diameter of ∼30 nm) (see **Figure S3a**, Supporting Information). This active opening mechanism was introduced into the MP to ensure a reliable opening of the structure also e.g. in the environment of a lipid membrane, where lateral membrane pressure could counteract a stable ‘fully open’ conformation. In order to controllably switch from the open to the intermediate state, all opening strands were designed with an additional 7-nt toehold region. This allows strand displacement of the opening strands upon the addition of fully complementary invader strands (‘anti-opening strands’), making the conformational switching of our MP fully reversible.

**Figure 1.**
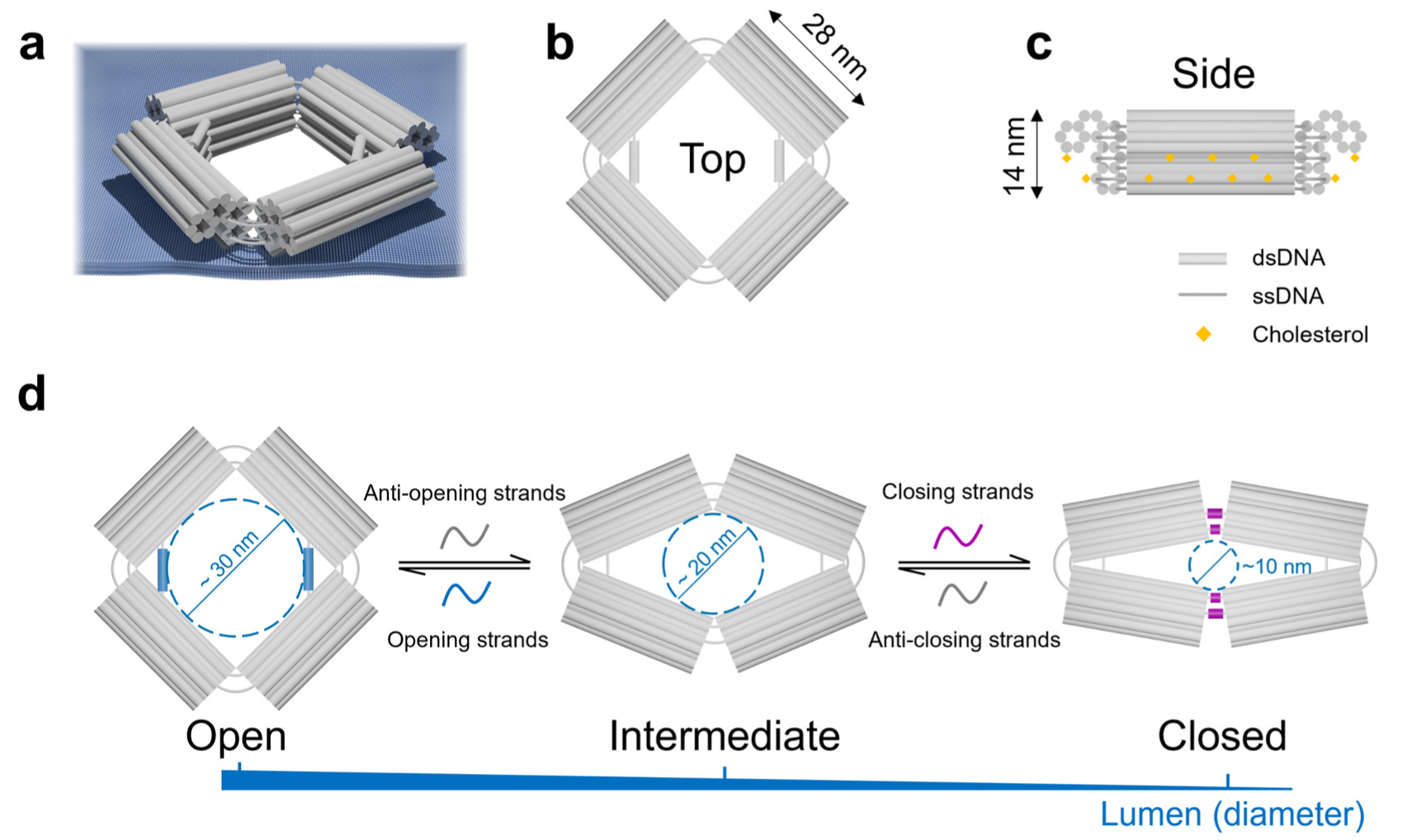
Design and working principle of the reconfigurable MechanoPore. (a) 3D illustration of the DNA nanopore in the open state (MP-O) when embedded in a lipid membrane. (b) Top and (c) side view of MP-O. Grey cylinders represent dsDNA, while the grey lines are ssDNA. Yellow diamonds represent schematically the attached cholesterol modifications. (d) Reversible conformational changes between three states (open, intermediate, and closed) of the MP in response to the addition of trigger strands (blue and magenta for opening and closing strands, respectively) and anti-trigger strands (grey).

Similarly, an active closing mechanism was implemented (**Figure S3c**, Supporting information). To achieve this, the MP was designed with flexible orthogonal ssDNA regions in two opposing corners. The addition of ‘closing strands’ forces the MP into its closed state with a small opening angle of about 20 - 25° and a significantly reduced inner lumen (diameter of ∼11 nm). Analogously to the active opening mechanisms, this conformational change into the closed state is fully reversible via the addition of the corresponding invader strands (‘anti-closing strands’) to displace the toehold-containing closing strands. Notably, the anti-closing strands and the opening strands can be added in one step to promote the immediate switching of the MP from its closed into its open state (and vice versa).

To enable the insertion of the MP into the hydrophobic interior of the lipid bilayer, 30 cholesterol-modified DNA strands were introduced, which protrude from the outer surface of the four bottom parts of the structure as well as from underneath the four caps.[33] The MP also features up to six handles for the attachment of Cy5-modified DNA strands for fluorescence imaging (see **Figure S1** for details of the positioning of the handles).

### Structural characterization

After the assembly of the MP, we initially assessed the quality of the folded structures by agarose gel electrophoresis (AGE) and negative-stain transmission electron microscopy (TEM), (Figure 2 and **Figure S4**, Supporting information). As can be seen from the respective images in Figure 2a-c, well-formed nanopores were obtained for the three different states. Their global shapes match well with the design of the three conformations, in which four rigid edges enclose the rhombus-shaped pore structure. The dsDNA struts stabilizing the open state can clearly be seen in the TEM micrographs as well as in the 2D classification images. The measured opening angles of the three states were determined to be approximately 90° (open), 50° (intermediate), and 25° (closed), respectively, as obtained from the TEM micrographs. The respective lumen sizes were determined to be 496 ± 160 nm^2^ (open), 350 ± 106 nm^2^ (intermediate), and 194 ± 144 nm^2^ (closed) (see Figure 2d). In principle, a variety of fixed opening angles of the MP can be achieved by tuning the lengths of the double-stranded regions and the single-stranded loops in the struts by choosing a different set of opening staples (see **Figures S3b** and **S5**, Supporting information).

**Figure 2.**
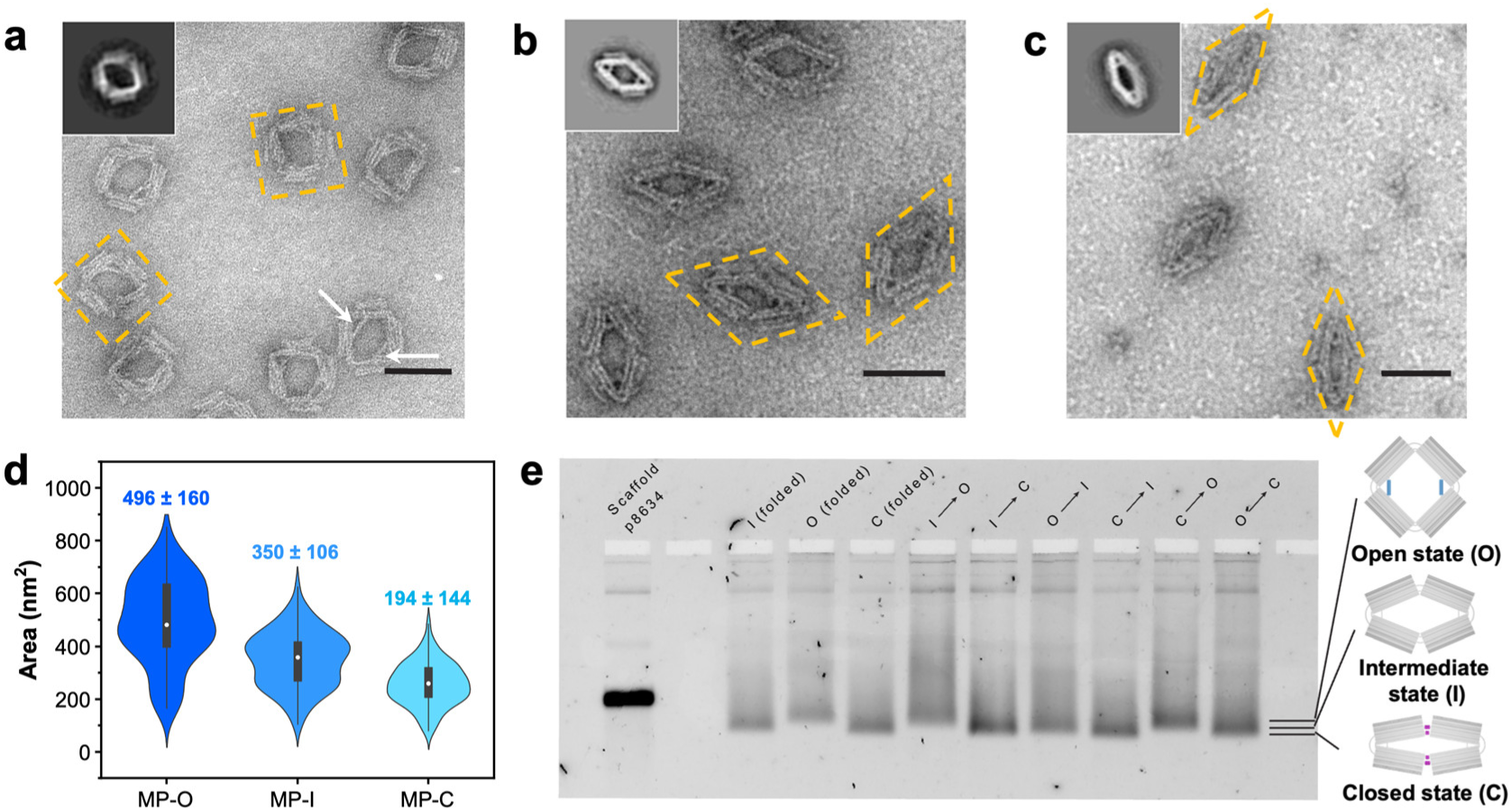
Negative-stain TEM images, 2D classification and AGE of the MP in its different configurations. (a-c) TEM images of the MP in its open, intermediate, and closed states. Insets: class averaging for 90 particles. Yellow dashed lines outline the shapes of representative structures in the TEM images. White arrows point to the double-stranded struts of the MP in the open state. Scale bars: 50 nm. (d) Statistical analyses of the inner area of MPs in different states. (e) Agarose gel of the MPs that were folded in different configurations or switched between different states in solution.

From the gel image in Figure 2e it can also be seen that the three states display slight differences in their electrophoretic mobility. Structures in the open state exhibit the slowest electrophoretic mobility, while structures in the closed state display the highest mobility. This can be easily explained by the effective overall volume increase from the closed to the open state. Although the intermediate state is an equilibrium between the open and the closed state, as it is not stabilized by external triggers, both the TEM analysis and AGE show that intermediate structures adopt a preferred conformation with an average opening angle of 50° and display an intermediate electrophoretic mobility between the open and the closed state.

In order to investigate the switching behaviour of the MPs, we performed 3D-DNA-PAINT[34] measurements as well as monitoring the electrophoretic mobility after the addition of the respective trigger strands. As can be seen in Figure 2e, shifts in electrophoretic mobility according to the structural change induced by the trigger strands can be observed as desired. To analyse the switching behavior in situ with 3D-DNA-PAINT, the MPs were anchored to a glass surface via neutravidin-biotin interactions (biotin handles located at the bottom part of only one of the four subunits; **Figure S2,** Supporting information). DNA-PAINT docking sites were positioned at the two opposing corners of the MPs without the struts, allowing for distance monitoring between the corners (see Figures 3 and **S2,** Supporting information). In the case of closed MPs, the distance between the binding sites might be smaller than the resolution of DNA-PAINT[37], thus leading to overlapping signals from the two corners of the MPs which could result in the erroneous exclusion of MPs with narrow opening angles from the data analysis. For reliable and unbiased distance measurements independent of the MP configuration, we employed a variant of DNA-PAINT called Resolution Enhancement by Sequential Imaging[35] (RESI). Labeling opposing corners with orthogonal imager binding sites and sequentially imaging them allows to unambiguously distinguish between the signals from the two corners.[36] Thus, the coordinates of the corners can be reliably identified and the distance between them can be precisely determined (see Methods and **Figure S7** for details about the image analysis).

**Figure 3.**
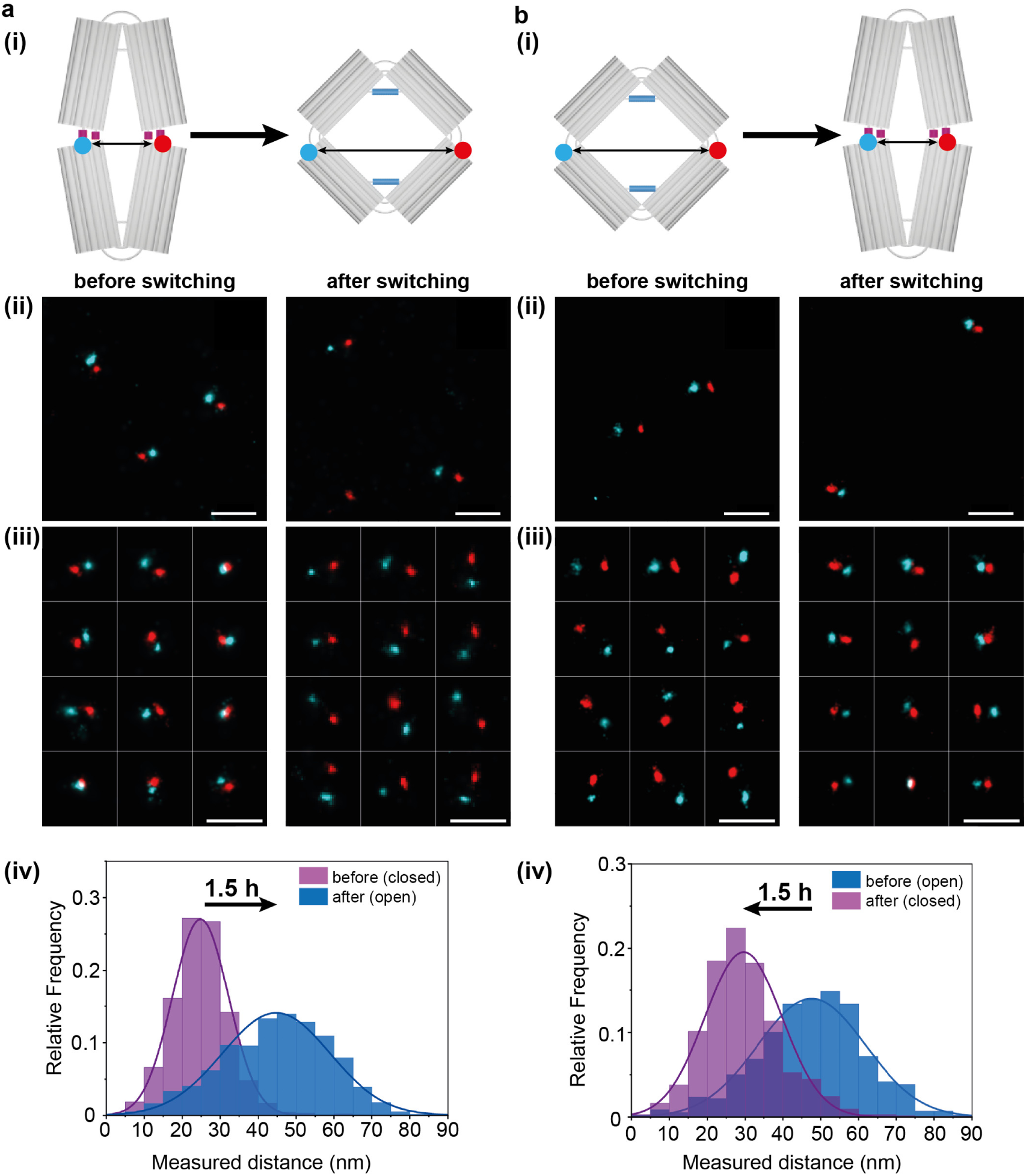
3D-DNA-PAINT measurements on MPs. (a) Conformational change from closed to open state with an incubation time of 90 min. (b) Conformational change from open to closed state after 90 min of incubation with the respective trigger and anti-trigger strands. Schematic representation of the switching of the MPs for the respective experiment (i), representative fields of view from all datasets (ii), overview over representative nanopores selected during data analysis, clearly showing the shift in the observed distances (iii) and the respective distance distributions for the switching experiments (iv). Scale bars: 100 nm.

We initially performed measurements on MPs pre-formed in the open, intermediate and closed states (see **Figure S6**, Supporting Information). The distance distributions obtained from the different nanopore configurations exhibit mean distances of 26.0 ± 8.4 nm (fit of a normal distribution, closed state), 36.2 ± 12.1 nm (intermediate state) and 47.5 ± 16.4 nm (open state) which match the expectations from the design and the characterization via TEM, taking into account the positions of the imager binding sites on the nanopore caps (Figure 2 and **Figure S4**, Supporting information). As a next step, we demonstrated *in situ* switching. We first performed a complete 3D-DNA-PAINT measurement on the MPs in their closed state. Then, the slide was thoroughly washed to remove all imager strands. After that, both the anti-closing strands and the opening strands were added in a twentyfold molar excess to the channel and the slide was incubated at 33 °C. In order to probe the time scale of the switching, we investigated two different incubation times (90 min and 3 h). After this, excess trigger strands were flushed out of the channel and a second full 3D-DNA-PAINT measurement was performed. Figure 3a shows the distance distributions for the MPs before and after the incubation with the trigger strands for 90 min, representative sections (fields of view, FOVs) from the aligned datasets and an overview of representative selected nanopores. The sample in the closed state exhibits a distance distribution with a peak centered at around 24.8 ± 7.4 nm which corresponds to an opening angle of around 20 - 24°, taking into account the positions of the handles on the MPs. After the switching procedure, the main peak in the distance distribution is shifted to 44.7 ± 14.2 nm, showing a successful conformational change to the open state. A similar behaviour could be observed after the longer incubation time of 3h (**Figure S7**, Supporting information), suggesting that 90 min is already sufficient. In both experiments, the obtained distance distributions closely correlate to those obtained for MPs pre-folded into the open state. To demonstrate full reversibility, we next tested the reverse switching of the MPs from the open into the closed state. Figure 3b shows representative FOVs from the data sets, examples for selected MPs and the corresponding distance distributions for the sample before and after incubation with the anti-opening and the closing strands at 33°C for 90 min. Again, we observed a pronounced shift of the mean of the distance distribution from around 47.6 ± 14.3 nm in the initial state to around 28.6 ± 10.2 nm, clearly demonstrating the successful switching to the closed state. These results confirm the robustness of the nanomechanical switching capabilities of the MPs.

### Reconstitution and permeability of MPs in lipid bilayers

After successful characterization of the MPs and confirmation of their controlled conformational switching *in situ,* we next tested their ability to insert into a lipid membrane. The insertion of large DNA origami nanopores into lipid membranes is far from trivial. As DNA molecules are hydrophilic, DNA nanostructures cannot penetrate the membrane spontaneously without lipidation by e.g. cholesterol modification. The number of cholesterol modifications required for successful insertion strongly depends on the size of the structure, and the incorporation of many cholesterol-modified staples can cause aggregation of the nanostructures.[33] However, careful tuning of the positioning of cholesterol modifications on the DNA origami (distance between adjacent modifications, handle lengths, etc.) can reduce the risk of aggregation. As can be seen in **Figure S8**, we do not observe significant aggregation or clustering of the nanostructures after cholesterol addition.

A further challenge is that large DNA nanostructures can adopt various orientations when inserted into the membrane and the incorporation yield of functional membrane spanning pores can be very low. While the embedding of small DNA origami nanopores into lipid membranes can often rely on spontaneous insertion into preformed lipid bilayers, the probability of insertion of large nanopores decreases dramatically with increasing size of the pores.[22] To address these challenges, we used the cDICE (continuous droplet interface crossing encapsulation) method to embed the nanoactuators into GUVs during liposome formation.[11, 37] This method involves injecting the DNA origami solution (with 2 MDa dextran) into a rotating chamber with a lipid-in-oil dispersion (inner layer) and a glucose solution (outer layer) (Figure 4a). The injected solution forms droplets in the oil phase and the GUVs form when the droplets cross the oil-water interface and can be collected from the outer solution after 30 min. The use of 2 MDa dextran in the DNA origami solution helps to drive the MPs to the lipid surface and increases their effective density, drastically enhancing their incorporation efficiency.[11] Due to its large size, the 2 MDa dextran remains in the GUV in the presence of MP (even in their open state).

**Figure 4.**
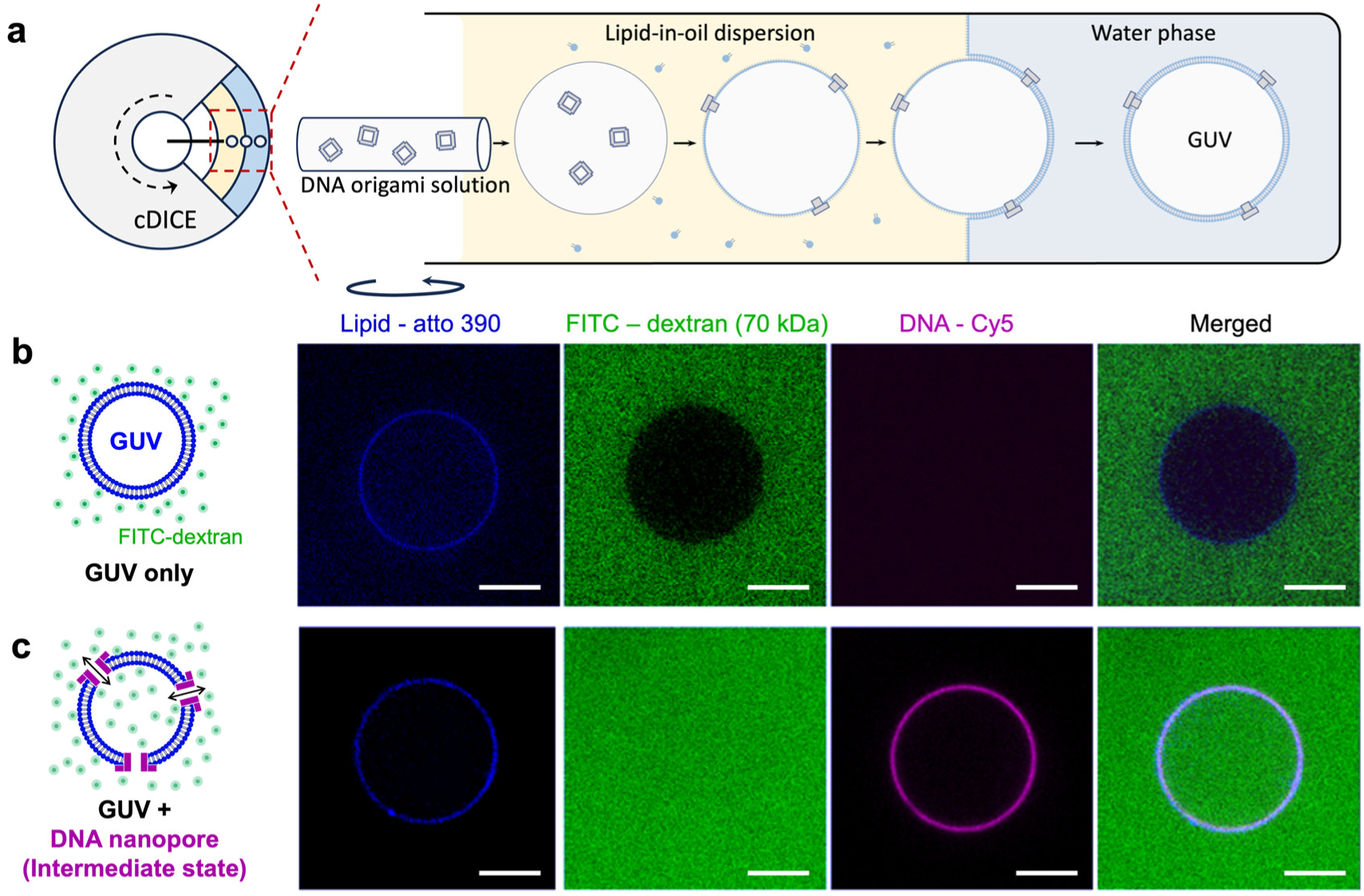
cDICE and permeability of the MPs reconstituted in GUVs. (a) Schematic representation of cDICE. (b) and (c) are the confocal images of GUV and GUV with the MPs (in the intermediate state), respectively. From left to right, pseudo-colours are blue = atto 390 (GUV), green = FITC (FITC-dextran), magenta = Cy5 (DNA), and the merged channel. Scale bars: 5 µm.

Initially, we used AFM to investigate the occurrence of insertion upon splashing GUVs onto a mica substrate, leading to a flat membrane (**Figure S9,** Supporting information). After MP insertion into the GUVs, we found some brighter spots on the membrane, which we attribute to the nanopores. A comparison of the heights of the bare lipid membrane, free MPs and the MPs located on the membrane (∼ 6, 12 and 5 nm respectively, whereby the last value is measured with respect to the height of the lipid membrane) strongly indicates that a part of the DNA origami structure is embedded into the membrane.

Since the MPs should function as size-selective transmembrane channels, we also investigated their reconstitution and pore-forming capabilities in GUVs using confocal microscopy. We introduced different fluorescence modifications into the lipid membrane of the GUVs (Atto 390, blue channel) and onto the MPs (Cy5, red channel) to visualize their presence and spatial distribution. FITC-modified dextran (molecular weight: 70 kDa, green channel) was used as cargo to test the permeability of the MPs.

In Figure 4, we initially analysed pure GUVs without any MPs. Confocal imaging clearly shows circular vesicles in the blue channel, while the fluorescence in the red channel is negligible, indicating the absence of MPs. By analyzing the fluorescence intensity in the green channel, we found that most of the GUVs exhibit a large intensity difference between inside and outside of the vesicles, suggesting that the FITC-dextran is unable to enter the GUVs. Quantitatively, we calculated the normalized fluorescence intensity difference *I_ndiff_ = (I_out_ – I_in_) / I_out_* of pure GUVs, which yields 0.21 ± 0.04 (error: S.D., N = 69) consistent with the results of unfilled GUVs from a previous study,[11] where *I_out_* and *I_in_* are the fluorescence intensities outside and inside the vesicles, respectively (**Figure S10**). This confirms that the lipid membrane blocks the translocation of the FITC-dextran and is therefore impermeable to such macromolecules. In the next step, MPs in their intermediate state were included in the GUV production with the cDICE technique. In contrast to the negative control, the images show the colocalization of Cy5 fluorescence with the fluorescence signal from the labeled lipids, clearly indicating the presence of MPs that are evenly distributed on the vesicle membrane. It should be noted, however, that the presence of a fluorescence signal does not necessarily imply complete insertion of structures into the lipid membranes. Heterogeneity in GUV samples[38] and the possibility that some DNA nanopores may only adhere to the membrane, not forming full transmembrane channels, resulted in a certain fraction of unfilled GUVs. We therefore analyzed the fluorescence intensity from the FITC-dextran, in order to determine if transmembrane channels were formed. The fluorescence intensity was found to be very similar inside and outside the GUVs (*I_ndiff_* ≈ 0, **Figure S10**, Supporting information). This suggests that the MPs can be successfully inserted into lipid bilayers, form transmembrane channels, and enable the transmembrane passage of macromolecules.

Subsequent FRAP assays were used to estimate the diffusion rate of dextran across the nanopores and the average distribution density of the nanopores on the GUV membrane (**Figure S11**, Supporting information). We compared the fluorescence recovery of pure GUVs and GUVs containing MPs, which were photobleached only in their interior after overnight incubation with 70 kDa FITC-dextran. As expected, an MP-containing GUV showed almost full recovery of fluorescence signal after a few minutes, while a GUV without MPs showed negligible recovery even with a longer time measurement (**Figure S11 b**-**e**). We then selected the MP-containing GUVs that showed almost equal intensity inside and outside (*I_ndiff_* ≈ 0) and employed the same FRAP assay to fit a diffusion model derived from Fick’s first law and estimated the number of nanopores (*N_p_*) in a GUV (**Figure S11f**). By analyzing the FRAP curves obtained from GUVs with different sizes, we found that the estimated number of pores *N_p_* per GUV varies from a few to several hundred.[11] For a 12 μm diameter vesicle, we estimated the number of functional pores to be *N_p_* = 125. While there is a large variation in *N_p_*, we do observe an overall increase in *N_p_* with increasing GUV size, roughly proportional to a cubic fit to the GUV diameter (**Figure S11g**), which is consistent with the assumption that MPs move from the inner GUV volume to the membrane during GUV formation.

### Size selectivity of the MechanoPores

Following the successful integration of MPs in GUVs, we set out to test the size-selective transport of macromolecules through MPs with different initial conformations. Here, 10, 70 and 150 kDa FITC-dextrans were used as molecular standards. The gyration diameter of dextrans with different MW can be estimated by the empirical equation *D_g_* = 0.072 × *MW*^0.48^ [39], resulting in values of approximately 6, 15, and 22 nm, respectively. Therefore, the transmembrane passage of large 150 kDa FITC-dextran should only be possible if successfully inserted MPs are present in their fully open state, whereas 10 kDa FITC-dextran should be able to pass through the nanopores even in their closed state. Extraction of the FITC fluorescence intensity inside and outside the boundary of GUVs allowed us to calculate the normalized intensity difference *I_ndiff_*. By comparing the different behaviours of pure GUVs and GUVs containing MPs, the histograms clearly indicate the existence of two types, corresponding to filled GUVs (*I_ndiff_*: *-*0.10 to 0.09) and unfilled GUVs (*I_ndiff_*: 0.09 to 0.33), (**Figure S10**, Supporting information). Thus, when *I_ndiff_* was lower than 0.09, the corresponding vesicle was marked as filled (see **Figure S12** for histograms of *I_ndiff_* for three independent checks of each state, Supporting information).

The experimental data displayed in Figure 5 shows the successful incorporation of all three different MPs into the GUV as indicated by their fluorescent signal in the membrane. In the open configuration, 10 kDa dextran (80 ± 9% filled), 70 kDa dextran (64± 8% filled), and 150 kDa dextran (46 ± 8% filled) could pass through the nanopores, as expected. MPs in the intermediate state only allowed the translocation of the 10 kDa dextran (86 ± 10% filled) and the 70 kDa dextran (55 ± 9% filled), while most of 150 kDa dextran (20 ± 9% filled) was blocked out. In the closed conformation, transmembrane passage ccould only be observed for the 10 kDa dextran (64 ± 7% filled). This indicates that, as designed, from closed to open conformations, the MPs show gradually increasing lumen sizes after insertion into the lipid membrane. This in turn translates into their ability to size-selectively translocate cargo across the lipid membranes or hinder molecules from entering the GUVs.

**Figure 5.**
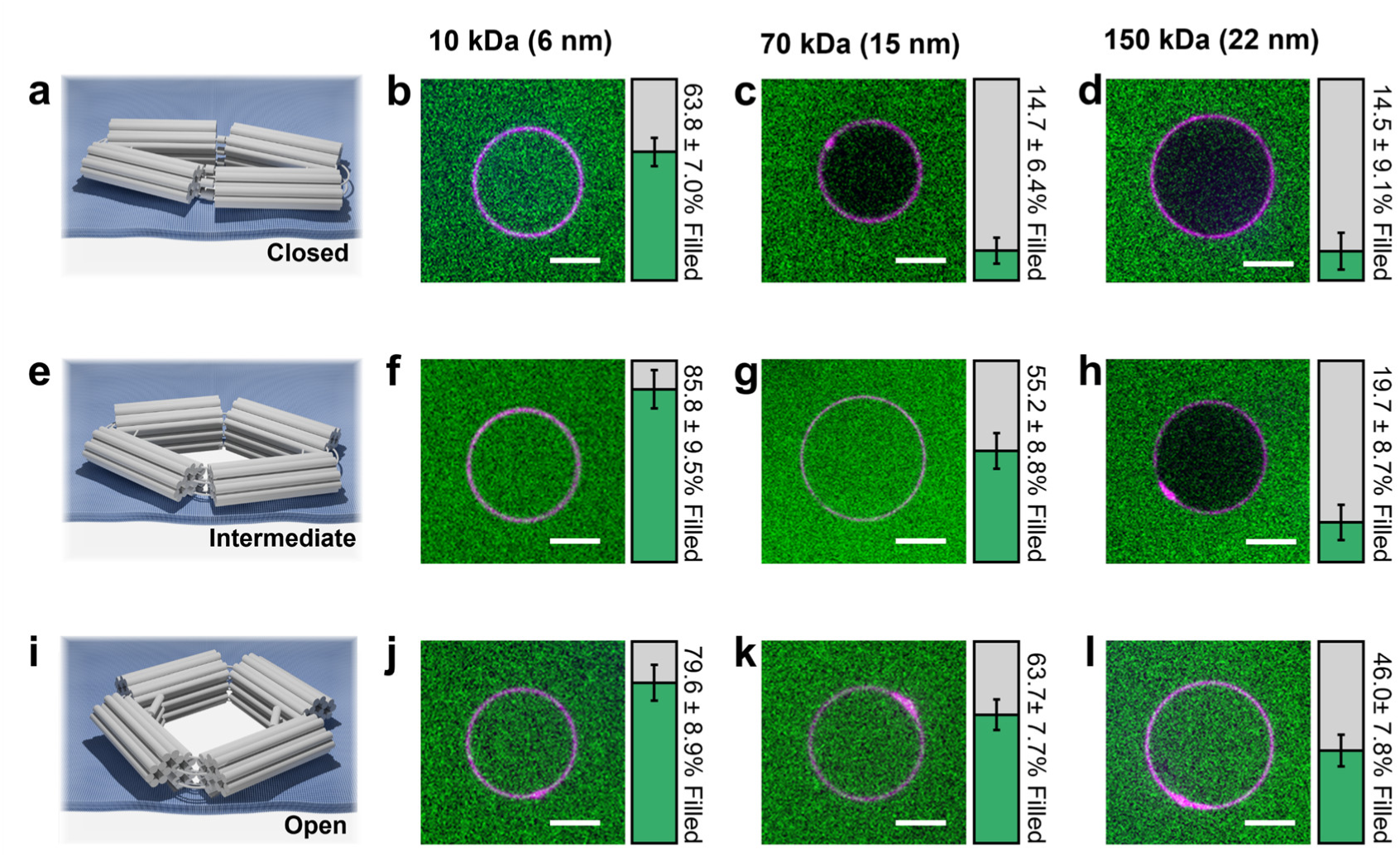
Size-selectivity of 3 different states of MPs. Confocal images of 10 kDa, 70 kDa and 150 kDa FITC-dextran influx in closed state (a-d), intermediate state (e-h), and open state (i-l) MPs. Each image shows the representative behaviour of more than 150 GUVs. Bar graphs on the right are the percentages of GUVs in which dextran influx is observed. Scale bars: 5µm. Error bars: S.D.; three independent experiments.

To further analyse the lumen sizes after incorporation into the lipid membrane, we used TEM analysis of splashed GUVs (**Figure S13**, Supporting information). Interestingly, the MPs exhibited a decreased average lumen size after membrane insertion (compared to values obtained in the absence of a lipid membrane) for all three configurations (O: from 496 ± 160 to 353 ± 110 nm^2^; I: from 350 ± 107 to 198 ± 73 nm^2^; C: from 194 ± 144 to 118 ± 52 nm^2^). This can be attributed to lateral pressure exerted by the lipid membrane, which has been reported to lead to deformations of DNA origami nanopores, reducing the actual lumen size of the nanostructures.[25] Interestingly, however, the intermediate state truly remained as such and did not get compressed to a lumen size close to the closed state. Although we cannot exclude partial deformation of the intermediate state, we can clearly distinguish between the closed and the intermediate state, suggesting that despite the semi-flexible nature of the structure itself, lateral membrane pressure is not strong enough to fully close the structure, and thus confirming the necessity for a controlled closing mechanism.

### In-situ reversible conformational changes of DNA nanoactuators in lipid bilayers

For future applications, it is of great importance that the MP can not only be switched between its different configurations in solution or when anchored to a surface, but also when embedded within a lipid membrane, having to exert forces against the lateral membrane pressure. In order to determine the ability of the MPs to reversibly change their conformation also after membrane insertion, we incorporated the MPs in their intermediate state into GUVs and subsequently added the respective trigger (and anti-trigger) strands followed by confocal imaging. Dye influx experiments were performed using the 70 kDa FITC-dextran as the standard cargo, as the translocation behaviour of this dextran allows for a clear identification of the nanopore conformation.

For this, a sample was divided into five batches that were triggered to undergo a different sequence of conformational changes: 1) intermediate → open, 2) intermediate → open → intermediate with a 2 h interval between each switch, 3) intermediate → closed, 4) intermediate → closed → intermediate with a 2 h interval between each switch, 5) adding buffer as a blank. For each sequence (1-5), we analysed part of the resulting sample by TEM after 2 h incubation at room temperature, while the remaining part was used for confocal imaging of dye influx with the 70 kDa FITC-dextran after 24 h. Importantly, both TEM analysis (**Figure S14**, Supporting information) as well as dye influx assays show that conformational switching to an open or closed state as well as reversible switching back to the intermediate state is still possible even after embedding into the lipid membrane.

As can be seen from Figure 6 (bar percentages are calculated from **Figure S15**, Supporting information), initially, MPs in the intermediate state showed an almost equal distribution of filled and unfilled GUVs (52 ± 7% filled). After adding the respective trigger strands, the percentage of filled GUVs changed to 66 ± 8% (intermediate → open) and 27 ± 7% (intermediate → closed), respectively. Structures switched from intermediate to open (or closed) and back to the intermediate state (i.e. processes 2 and 4) displayed a similar distribution and percentage of filled GUVs (56 ± 7% and 53 ± 9% filled) as the non-actuated MPs, indicating the achievement of reversibly controllable conformational changes. Interestingly, the incorporation of pre-formed closed MPs showed a slightly lower percentage of filled GUVs (cf. Figure 5c). We attribute this to a partially sterically hindered position of the closing mechanism after insertion into the lipid membrane. Since the linking regions of the corners (and thereby the regions that are responsible for the active closing mechanism) are located on the outside of the MP, the diffusion of the closing strands to their respective binding sites may be partially obstructed. Nevertheless, after the addition of anti-opening strands, a clear difference between the closed and the intermediate state can be observed.

**Figure 6.**
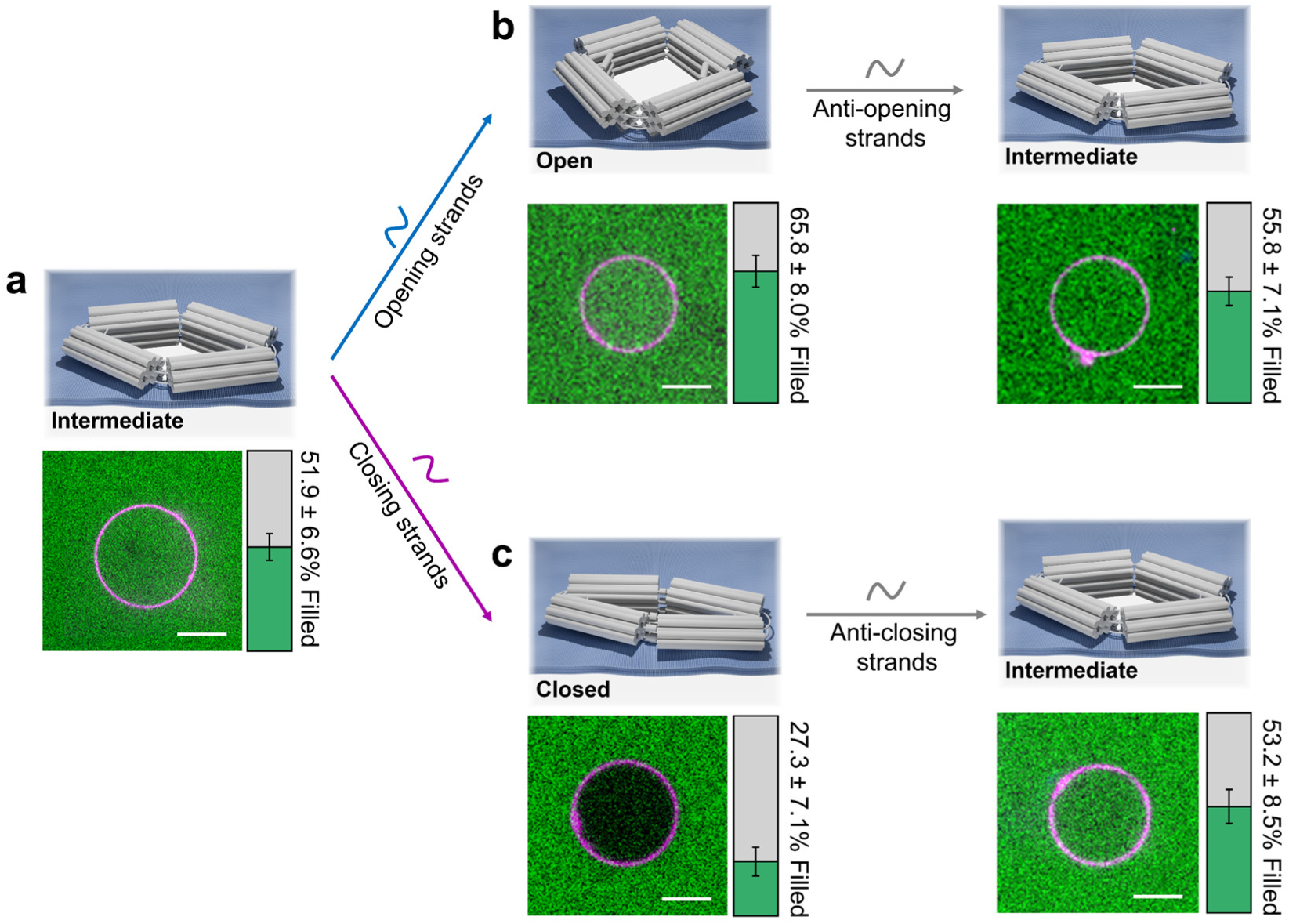
Reversible conformational changes of MPs embedded in GUVs and their permeability. Schematic representation (top) and confocal image (bottom) of (a) the intermediate state with 70kDa FITC-dextran, (b) i → o → i switching, when adding opening and subsequently anti-opening strands, followed by 70 kDa FITC-dextran addition, and (c) i → c → i switching, when adding the closing and subsequently anti-closing strands followed by 70 kDa FITC-dextran addition. The distribution and percentage of filled GUVs in the dye influx experiments is shown in the respective colour bars. Green: filled, and grey: unfilled. Error bars: S.D. from three independent samples measured. Scale bars: 5 µm. N.B.: in all cases dextrans were added *after* addition of the respective trigger strands.

Overall, these experiments successfully show that active conformational switching and reversible control over the lumen size of DNA origami nanopores can be achieved with at least three distinct and stable states.

## Conclusions

Here, we present a dynamic, three-state reconfigurable DNA origami MechanoPore that exhibits size-selective transport of biomolecules across lipid membranes on-demand. We overcome the challenge associated with reconstitution of the large nanopores (edge length of around 28 nm and ∼500 nm^2^ lumen) in lipid bilayers by using the cDICE technique for direct integration of the MPs in GUVs.

The MPs are designed as nanoactuators that can be robustly switched between three distinct conformations in a well-controlled and fully reversible manner via an external trigger mechanism (addition and displacement of DNA strands). The resulting lumen dimensions can thus be tuned from ∼100 to 500 nm^2^. The rationally engineered opening and closing mechanisms actively promote the conformational switching of the MP not only in solution, but also when anchored to a surface and, most importantly, after insertion into lipid membranes.

To the best of our knowledge, this is the first experimental demonstration of reversible switching of a DNA origami transmembrane channel in lipid membranes with multiple well-defined structural states. We confirm the nanomechanical switching at individual nanopore level by accurately monitoring the states of the MPs via 3D-DNA-PAINT, which enables the distinction between open, intermediate and closed configurations. We additionally successfully demonstrate reversible nanomechanical cycling in GUVs using confocal imaging of dye influx assays.

The various stable configurations (open, intermediate, closed) within the lipid bilayer result in vastly different enclosed lumen sizes, allowing for the passage/transport of large molecules in the open state of the nanopores, yet also enabling the effective exclusion of small molecules from transmembrane passage if the very same nanopores are switched to their closed state. By employing FITC-dextrans of various sizes ranging from 6 to 22 nm in diameter, we clearly observe size-selective transport of molecules, with higher percentages of GUV filling for the most open MP configuration, and conversely blocked transport for smaller pore diameters (apart from the smallest measured 10 kDa dextrans). We note that the MP design is highly versatile, enabling the generation of nanopores with a range of stable lumen dimensions by simply changing the length of the triggers. We thus expect that this approach can be readily extended to molecular sieving applications that require even finer control of translocation sizes.

Our results demonstrate the potential of structurally adaptable DNA origami nanopores for applications in molecular delivery, sorting and biosensing. We anticipate that our findings will pave the way towards more advanced dynamic nanodevices with potential uses in the field of controlled drug delivery and molecular diagnostics, as well as biomimetics and synthetic cell research, where controlled transport of biological macromolecules through large, stable channels is crucial. We thus highlight the wide applicability of these nanopores for both fundamental and applied biophysics research.

## Methods

### DNA origami folding and purification

The DNA origami nanoactuator was folded using 15 nM of the scaffold p8634, 100 nM of each staple strand in buffer containing 5 mM Tris, 1 mM EDTA (pH = 8) and 20 mM MgCl_2_. The mixture was heated to 65 °C and held at this temperature for 5 min, then cooled down to 20 °C over a period of 20 hours. All additional handle staples were incorporated during folding (see **Figures S1** and **S2** and **Table S1** for handle positions and sequences).

Purification of the folded DNA origami nanoactuators from excess staple strands was achieved using ultrafiltration with 0.5 mL Amicon centrifugal filter units with 100 kDa molecular weight cut-off (Merck Millipore). The filter units were pre-washed with buffer (1x TAE, 5 mM MgCl_2_), then the folding solution was loaded into the filters and centrifuged at 5000 rcf for 6-8 min. This step was repeated at least five times with fresh buffer (1x TAE, 5 mM MgCl_2_) added in every centrifugation round.

Switching of the nanoactuators in solution was carried out by incubating the folded DNA origami with a 10-fold molar excess of the respective trigger strands at 33 °C for 3 hours.

### Transmission electron microscopy (TEM) imaging

After the DNA origami nanostructures were diluted to 10 nM concentration, 10 µl of DNA origami solution were incubated for 2 min on a carbon-coated copper grid (Plano GmbH, Formvar/carbon 300 mesh) that had been plasma-cleaned for 30 s. For nanopore-containing GUV samples, additional 2 min incubation time is required. The solution was then removed with the help of filter paper and the grid was stained with 5 µl of 2% uranyl formate solution for 45 s. Images were obtained on a Jeol-JEM-1230 TEM operating at an acceleration voltage of 80 kV. RELION 3 (https://relion.readthedocs.io/) was used to do the 2D classification analysis.

### Atomic force microscopy (AFM) measurements of nanopore-containing GUV samples

AFM imaging was performed with a NanoWizard 4 (JPK Instrument AG) in QI^TM^ mode in 1 x TAE buffer with 10 mM MgCl_2_ using SCANASYST-FLUID+ probe (Bruker, Si N-type; L: 70 µm, W: 10 µm, T: 2 µm; Spring constant: 0.7 N/m). 5 µL of sample was deposited on the fresh cleaved mica for 15 min, rinsed with 200 µL 1 x TAE buffer with 10 mM MgCl_2_ two times. Image acquisition: with 256 x 256 pixel resolution.

### Agarose gel electrophoresis

The DNA origami samples were diluted to 10 nM concentration in 1 x TAE buffer containing 5 mM MgCl_2_ and then mixed with loading dye containing Orange G and glycerol.

Subsequently, the samples were loaded onto a 2% agarose gel that was pre-stained with 0.01% Sybr Safe. The gel was run on ice for 150 min at 75 V (running buffer: 1 x TAE with 11 mM MgCl_2_). Gel imaging was then carried out on a Typhoon FLA-9000.

### DNA-PAINT experiments

#### Microscope setup

Fluorescence imaging with TIRF illumination was carried out on an inverted microscope (Nikon Instruments, Eclipse Ti2) with the Perfect Focus System, applying an objective-type TIRF configuration equipped with an oil-immersion objective (Nikon Instruments, Apo SR TIRF x100, NA 1.49, Oil). A 560-nm laser (MPB Communications, 1 W) was used for excitation. The laser beam was passed through a cleanup filter (Chroma Technology, ZET561/10) and coupled into the microscope objective using a beam splitter (Chroma Technology, ZT561rdc). Fluorescence was spectrally filtered with an emission filter (Chroma Technology, ET600/50m and ET575lp) and imaged on an sCMOS camera (Andor, Zyla 4.2 Plus) without further magnification, resulting in an effective pixel size of 130 nm (after 2 x 2 binning). The readout rate was set to 200 MHz. Images were acquired by choosing a region of interest with a size of 512 x 512 pixels. 3D imaging was performed using a cylindrical lens (Nikon Instruments, N-STORM) in the detection path. Raw microscopy data was acquired using μManager[40] (Version 2.0.1).

#### Buffers

The following buffers were used for sample preparation and imaging:

- Buffer A: 10 mM Tris pH 8, 100 mM NaCl and 0.05% Tween-20
- Buffer B: 10 mM MgCl_2_, 5 mM Tris-HCl pH 8, 1 mM EDTA and 0.05% Tween-20, pH 8
- Buffer B20: 20 mM MgCl_2_, 5 mM Tris-HCl pH 8, 1 mM EDTA and 0.05% Tween-20, pH 8
- 100x Trolox was made by adding 100 mg Trolox to 430 μl of 100% methanol and 345 μl of 1 M NaOH in 3.2 ml water.
- Imager solution: Buffer B20, 1x Trolox, pH 8, supplemented with Cy3B-coupled DNA imager strands.
- 20nm grid DNA origami folding buffer: 10 mM Tris, 1 mM EDTA, 12.5 mM MgCl_2_, pH 8
- FoB5 Buffer: 5 mM Tris, 1 mM EDTA, 5 mM MgCl_2_, 5 mM NaCl, pH 8.5

#### 20 nm grid DNA origami self-assembly and purification

The 20 nm DNA origami grid structures were designed with the Picasso design tool.[34] We followed the design used in Reinhardt et al.[35] and extended the 3’ end of staple strands at the grid nodes with the docking strand sequence 7xR3 (CTCTCTCTCTCTCTCTCTC). Self-assembly of DNA origami was accomplished in a one-pot reaction mix with a total volume of 40 μl, consisting of 10 nM scaffold strand, 100 nM folding staples, 250 nM biotinylated staples and 1 μM staple strands with docking site extensions in folding buffer. The reaction mix was then subjected to a thermal annealing ramp using a thermocycler. Thereby, the reaction mix was first incubated at 80 °C for 5 min, cooled using a temperature gradient from 80 to 65 °C in steps of 1 °C per 30 s and subsequent steps from 65 to 20 °C in steps of 1 °C per 1 min. The mix was finally incubated at 20 °C for 5 min and finally held at 4 °C.

The 20 nm grids were purified via ultrafiltration using Amicon Ultra centrifugal filters with a 50 kDa molecular weight cutoff (50 kDa MWCO; Merck Millipore, cat: UF505096). The filter units were first equilibrated with 500 μL of FoB5 buffer and centrifuged at 10,000g for 5 min. Then, the folded DNA origami were brought to 500 μL with FoB5 buffer, added to the filters, and centrifuged for 3.5 min at 10,000g. This process was repeated twice. Purified DNA origami were recovered into a new tube by centrifugation for 5 min at 5,000g.

#### DNA origami sample preparation and imaging

The samples for DNA-PAINT measurements were prepared in a six-channel ibidi slide. First, 80 µl of BSA-biotin (1 mg/ml, dissolved in buffer A) were incubated in each channel for 5 min, followed by washing of the channels with 450 µl buffer A. In the next step, 100 µl of neutravidin (0.5 mg/ml, dissolved in buffer A) were flushed into the channels and allowed to bind for 5 min. After washing with 150 µl of buffer A and subsequently with 450 µl of buffer B, gold nanoparticles (90 nm, diluted 1:19 in buffer B) were flushed into the channels and incubated for 5 min. The subsequent washing with 450 µl buffer B was followed by the addition of 80 µl of a DNA origami grid structure (∼ 200 pM) (serving for drift correction and alignment of RESI rounds) and incubation for 5 min. Then, the channels were again flushed with 450 µl of buffer B and the DNA origami nanopores equipped with biotin and handles for the DNA-PAINT imager strands (∼ 500 pM) were incubated in the channels for 5 min, followed again by washing with 450 µl of buffer B.

Finally, 180 μl of the imager solution supplemented with 300 to 500 pM R4 imager (GTGTGT-Cy3B) and 100 to 150 pM R3 imager (GAGAGAG-Cy3B) was flushed into the chamber. The chamber remained filled with imager solution and the imaging was performed. In between imaging rounds, the sample was washed with buffer B20 until no residual signal from the previous imager solution was detected. Then, the next imager solution supplemented with 1 to 1.5 nM R6 imager and 125 to 150 pM R3 imager was introduced and the second RESI round was imaged. For each round, 30000 frames were acquired with an exposure time of 100 ms and a laser power of 40 mW at the objective.

#### DNA-PAINT analysis

Raw fluorescence data were subjected to super-resolution reconstruction using the Picasso software package[34] (latest version available at https://github.com/jungmannlab/picasso). Drift correction was performed sequentially with gold particles and single DNA-PAINT docking sites serving as fiducials. Alignment of subsequent imaging rounds was performed iteratively, starting with a redundant cross-correlation and followed by fiducial alignment using 20 nm grids.

#### Distance measurements

The aligned data of the first (R3 & R4) and the second (R3 & R6) round were opened in Picasso Render. Regions of interest containing single nanopores with signal from both the R4 and the R6 site were manually selected in Picasso for further analysis. RESI analysis was performed by applying Density-Based Spatial Clustering of Applications with Noise (DBSCAN)[41] for each round separately, yielding groups of localizations, where each group originates from an individual binding site. Only regions where exactly one R4 and exactly one R6 group were detected were kept for further analysis. RESI localizations were calculated as previously described[35] as weighted means of the DNA-PAINT localization groups achieving an average precision of 0.76 nm. Finally, the Euclidian distance between the R4 and R6 site, representing the opening distance of the pore, was calculated within each region of interest.

### Fluorescence experiments

#### cDICE buffers and solutions

The following buffers were used for cDICE preparation:

- Inner solution: 50 mM Tris-HCl pH 7.4, 5 mM MgCl_2_, 15 μM 2 MDa dextran, 3 nM MechanoPores.
- Outer solution: ∼135 mM glucose in Milli-Q water, osmotically matched to the inner solution.
- Oil phase: mixture of silicone and mineral oil (4:1 ratio), lipids 0.2 mg/mL.

#### Preparation of lipid-in-Oil suspension

The lipid-in-oil suspension was prepared as in Van de Cauter et al.[37] and used immediately. DOPC and DOPE-Atto390 lipids solubilized in chloroform were mixed at 99.9:0.1 ratio (0.2 mg/mL) and blow-dried with pure nitrogen. Inside a glovebox filled with pure nitrogen, lipids were subsequently resolubilized with anhydrous chloroform. The freshly prepared mixture of silicone and mineral oil in 4:1 ratio was added to the lipids dropwise while vortexing at 1400 rpm. The lipid-in-oil solution was finally vortexed at 2800 rpm for 2 min and further sonicated in an ice bath for at least 15 min.

#### Imaging wells and passivation

The cover glass (24 x 60 mm) was first rinsed by ethanol and water and dried out by nitrogen flow. The bottom part of 1000 μL white pipette tips was cut and glued on the cover glass to serve as imaging wells. 200 µL BSA (5 mg/mL) was added to each imaging cell and incubate at room template for at least 30 min. BSA was gently removed from each well and the wells were rinsed with 200 μL outer buffer before use.

#### cDICE equipment and GUV production

The detailed protocol for GUV production was reported in Van de Cauter et al.[37] Briefly, cDICE chamber was first set to rotate at 300 rpm. 500 μL outer solution was pipetted into the rotating chamber and subsequently 6 mL of lipid-in-oil solution was added. The two phases are stacked as illustrated in Figure 4a. The tip of a 100 μm PEEK capillary tube (211633-3, BGB) was inserted into the oil phase of the rotating chamber, allowing for a continuous supply of inner aqueous solution into the oil phase by a syringe pump (Harvard Apparatus) set at 20 μL/min. After all the solution was injected into the chamber, the rotation speed (voltage) was gently turned down. The chamber was slightly tilted, and the oil phase was then removed as much as possible (4 to 5 mL). The chamber was set aside, tiled so that GUVs could sediment to the bottom of chamber. After 30 min, GUVs were collected from the outer aqueous solution with a 200 uL pipette tip (cut off the shape tip) and moved to a pre-passivated imaging well to carry out the fluorescence experiments.

#### Data collection and analysis

Fluorescence images were acquired using spinning disk confocal laser microscopy (Nikon Eclipse Ti-2, 100× objective, Photometrics Kinetix sCMOS) Nikon NIS software. FITC-dextran (10 kDa, 70 kDa, or 150 kDa) was gently mixed with GUVs to the final concentration of 2 μM. After 24 h incubation at room temperature, fluorescence was measured with 405 nm, 488 nm and 640 nm excitation.

#### Reversible conformational changes and influx

A batch of MP-I containing GUVs was divided into 5 samples with intermediate-state DNA nanoactuators as the baseline and each of the five batches was triggered to undergo a different set of conformational changes: 1) adding opening strand, 2) adding opening strand and anti-opening strand after a 2 h interval, 3) adding closing strand, 4) adding closing strand and anti-closing strand after a 2 h interval 5) adding buffer as blank. After that, FITC-dextran (70 kDa) was gently mixed with GUVs to the final concentration of 2 μM. After 24 h incubation at room temperature, the fluorescence was measured with 405 nm, 488 nm and 640 nm excitation.

Fluorescence images were analysed and processed using ImageJ (v1.54d).

#### Fluorescence recovery after photobleaching (FRAP) assay and diffusion model

To photobleach the FITC-dextran, we employed scanning with a 488 nm laser (at 9.8 mW) over the region of interest. To measure the recovery signal, frames were collected every 15 s, starting right after the photobleaching event.

#### Diffusion model

The details of diffusion model for nanopore were reported in Fragasso et al.[11] Briefly, the FRAP data were transferred to normalized intensity difference *I_ndiff_ = (I_out_ – I_in_) / I_out_,* and fitted with a diffusion model derived from Fick’s law of diffusion:

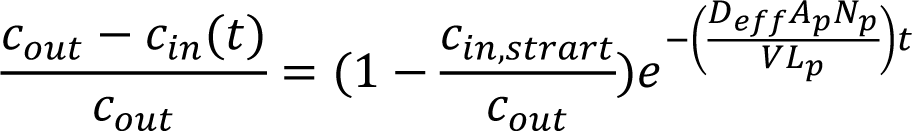

where *c_in,start_/c_out_* and *N_p_* were free parameters of the fit, all other parameters and constant are known or can be calculated. In particular, the area of pore *A_p_* = 197.8 nm^2^ was obtained from TEM experiments (**Figure S13**). *V* is the volume of the GUV, which can be calculated from the diameter obtained from confocal images. The channel length *L_p_* = 14 nm is the height of the nanopore. The effective diffusion constant *D_eff_* = 0.1 μm^2^/s of 70 kDa dextran was calculated using the formula from Dechadilok and Deen[42]:

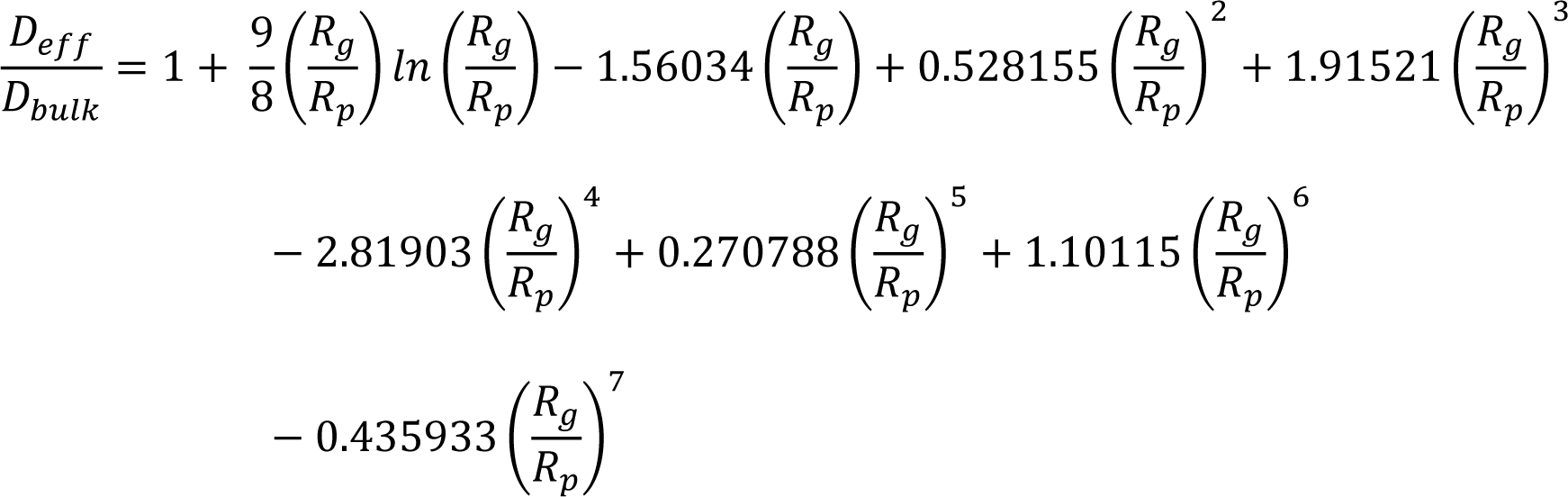

Where *R_g_* is the gyration radius of molecule (for 70 kDa dextran, R_g_ = 7.5 nm), *R_p_* is the pore radius (here 10 nm), and *D_bulk_* is the diffusion constant of the molecule in free solution (for 70 kDa dextran, *D_bulk_* = 45.87 μm^2^/s as we were using low concentration).[43]

## Supporting Information

Supporting Information is available from the Wiley Online Library or from the authors.

## Data availability

The data that support the findings of this study are available in the supplementary material of this article.

## Acknowledgements

Z.Y. was supported by an NWO-ENW-XS grant (Project MechanoPore). S.C. was supported by the ERC starting grant (SIMPHONICS, No. 101041486) and a Delft Technology Fellowship. A.H-J acknowledges financial support from the German Research Foundation (DFG) through SFB1032 (Nanoagents) project A06 and the Emmy Noether program (project no. 427981116). C.D. acknowledges financial support by the NWO program OCENW.GROOT.2019.068, ERC Advanced Grant no. 883684, and the NanoFront and BaSyC programs of NWO/OCW. R.J. and S.C.M.R. acknowledge support by the German Research Foundation through SFB1032 (project A11, no. 201269156). A.B. and S.C.M.R. acknowledge support by the IMPRS-LS graduate school. We thank Marianne Braun and Ursula Weber for assistance with TEM imaging.

## References

[1] S. Howorka, Nat. Nanotechnol. 2017, 12, 619.

[2] A. Dorey, S. Howorka, Nat. Chem. 2024, 16, 314.

[3] I. C. Nova, J. Ritmejeris, H. Brinkerhoff, T. J. R. Koenig, J. H. Gundlach, C. Dekker, Nat. Biotechnol. 2023.

[4] Y.-L. Ying, Z.-L. Hu, S. Zhang, Y. Qing, A. Fragasso, G. Maglia, A. Meller, H. Bayley, C. Dekker, Y.-T. Long, Nat. Nanotechnol. 2022, 17, 1136.

[5] P. D. E. Fisher, Q. Shen, B. Akpinar, L. K. Davis, K. K. H. Chung, D. Baddeley, A. Saric, T. J. Melia, B. W. Hoogenboom, C. Lin, C. P. Lusk, ACS Nano 2018, 12, 1508.

[6] C. Dekker, Nat. Nanotechnol. 2007, 2, 209.

[7] K. Shimizu, B. Mijiddorj, M. Usami, I. Mizoguchi, S. Yoshida, S. Akayama, Y. Hamada, A. Ohyama, K. Usui, I. Kawamura, R. Kawano, Nat Nanotechnol 2022, 17, 67.

[8] Y. Xing, A. Rottensteiner, J. Ciccone, S. Howorka, Angew. Chem. Int. Ed. 2023, 62, e202303103.

[9] Y. Xing, A. Dorey, L. Jayasinghe, S. Howorka, Nat. Nanotechnol. 2022, 17, 708.

[10] Q. Shen, Q. Xiong, K. Zhou, Q. Feng, L. Liu, T. Tian, C. Wu, Y. Xiong, T. J. Melia, C. P. Lusk, C. Lin, J. Am. Chem. Soc. 2022, 145, 1292.

[11] A. Fragasso, N. De Franceschi, P. Stömmer, E. O. van der Sluis, H. Dietz, C. Dekker, ACS Nano 2021, 15, 12768.

[12] S. Dey, A. Dorey, L. Abraham, Y. Xing, I. Zhang, F. Zhang, S. Howorka, H. Yan, Nat. Comm. 2022, 13, 2271.

[13] P. W. K. Rothemund, Nature 2006, 440, 297.

[14] S. M. Douglas, H. Dietz, T. Liedl, B. Högberg, F. Graf, W. M. Shih, Nature 2009, 459, 414.

[15] B. Shen, P. Piskunen, S. Nummelin, Q. Liu, M. A. Kostiainen, V. Linko, ACS Appl. Mater. Interfaces 2020, 3, 5606.

[16] C. Lanphere, J. Ciccone, A. Dorey, N. Hagleitner-Ertuğrul, D. Knyazev, S. Haider, S. Howorka, J. Am. Chem. Soc. 2022, 144, 4333.

[17] M. Langecker, V. Arnaut, T. G. Martin, J. List, S. Renner, M. Mayer, H. Dietz, F. C. Simmel, Science 2012, 338, 932.

[18] S. Iwabuchi, I. Kawamata, S. Murata, S.-i. M. Nomura, Chem. Commun. 2021, 57, 2990.

[19] S. Hernández-Ainsa, U. F. Keyser, Nanoscale 2014, 6, 14121.

[20] K. Göpfrich, A. Ohmann, U. F. Keyser, in Nanopore Technology, Vol. 2186 (Ed: M. A. V. Fahie), Springer US, New York, NY 2021.

[21] K. Göpfrich, C.-Y. Li, M. Ricci, S. P. Bhamidimarri, J. Yoo, B. Gyenes, A. Ohmann, M. Winterhalter, A. Aksimentiev, U. F. Keyser, ACS Nano 2016, 10, 8207.

[22] R. P. Thomsen, M. G. Malle, A. H. Okholm, S. Krishnan, S. S.-R. Bohr, R. S. Sørensen, O. Ries, S. Vogel, F. C. Simmel, N. S. Hatzakis, J. Kjems, Nat. Comm. 2019, 10, 5655.

[23] T. Diederichs, G. Pugh, A. Dorey, Y. Xing, J. R. Burns, Q. Hung Nguyen, M. Tornow, R. Tampé, S. Howorka, Nat. Comm. 2019, 10, 5018.

[24] J. R. Burns, A. Seifert, N. Fertig, S. Howorka, Nat. Nanotechnol. 2016, 11, 152.

[25] Y. Xing, A. Dorey, S. Howorka, Adv. Mater. 2023, 35, 2300589.

[26] J. J. Funke, H. Dietz, Nat. Nanotechnol. 2015, 11, 47.

[27] L. Zhou, A. E. Marras, H.-J. Su, C. E. Castro, ACS Nano 2013, 8, 27.

[28] A. Kucinic, C. M. Huang, J. Wang, H. J. Su, C. E. Castro, Nanoscale 2023, 15, 562.

[29] A. E. Marras, L. Zhou, H.-J. Su, C. E. Castro, Proc. Natl. Acad. Sci. 2015, 112, 713.

[30] M. Centola, E. Poppleton, S. Ray, M. Centola, R. Welty, J. Valero, N. G. Walter, P. Šulc, M. Famulok, Nat. Nanotechnol. 2023, 19, 226.

[31] Y. Ke, T. Meyer, W. M. Shih, G. Bellot, Nat. Comm. 2016, 7, 10935.

[32] S. M. Douglas, A. H. Marblestone, S. Teerapittayanon, A. Vazquez, G. M. Church, W. M. Shih, Nucleic Acids Res. 2009, 37, 5001.

[33] J. K. Daljit Singh, M. T. Luu, J. F. Berengut, A. Abbas, M. A. B. Baker, S. F. J. Wickham, Membranes 2021, 11, 950.

[34] J. Schnitzbauer, M. T. Strauss, T. Schlichthaerle, F. Schueder, R. Jungmann, Nat. Protoc. 2017, 12, 1198.

[35] S. C. M. Reinhardt, L. A. Masullo, I. Baudrexel, P. R. Steen, R. Kowalewski, A. S. Eklund, S. Strauss, E. M. Unterauer, T. Schlichthaerle, M. T. Strauss, C. Klein, R. Jungmann, Nature 2023, 617, 711.

[36] R. Jungmann, M. S. Avendaño, J. B. Woehrstein, M. Dai, W. M. Shih, P. Yin, Nat. Methods 2014, 11, 313.

[37] L. Van de Cauter, F. Fanalista, L. van Buren, N. De Franceschi, E. Godino, S. Bouw, C. Danelon, C. Dekker, G. H. Koenderink, K. A. Ganzinger, ACS Synth. Biol. 2021, 10, 1690.

[38] E. Baykal-Caglar, E. Hassan-Zadeh, B. Saremi, J. Huang, Biochim. Biophys. Acta, Biomembr. 2012, 1818, 2598.

[39] R. Hanselmann, W. Burchard, R. Lemmes, D. Schwengers, Macromol. Chem. Phys. 1995, 196, 2259.

[40] A. D. Edelstein, M. A. Tsuchida, N. Amodaj, H. Pinkard, R. D. Vale, N. Stuurman, J. Biol. Methods 2014, 1, e10.

[41] M. Ester, H.-P. Kriegel, X. Xu, In Kdd 1996, 96, 226.

42. P. Dechadilok, W. M. Deen, Ind. Eng. Chem. Res. 2006, 45, 6953.

[43] M. B. Albro, V. Rajan, R. Li, C. T. Hung, G. A. Ateshian, Cell. Mol. Bioeng. 2009, 2, 295.

