## Supporting information for "Compliant DNA Origami Nanoactuators as Size-Selective Nanopores"

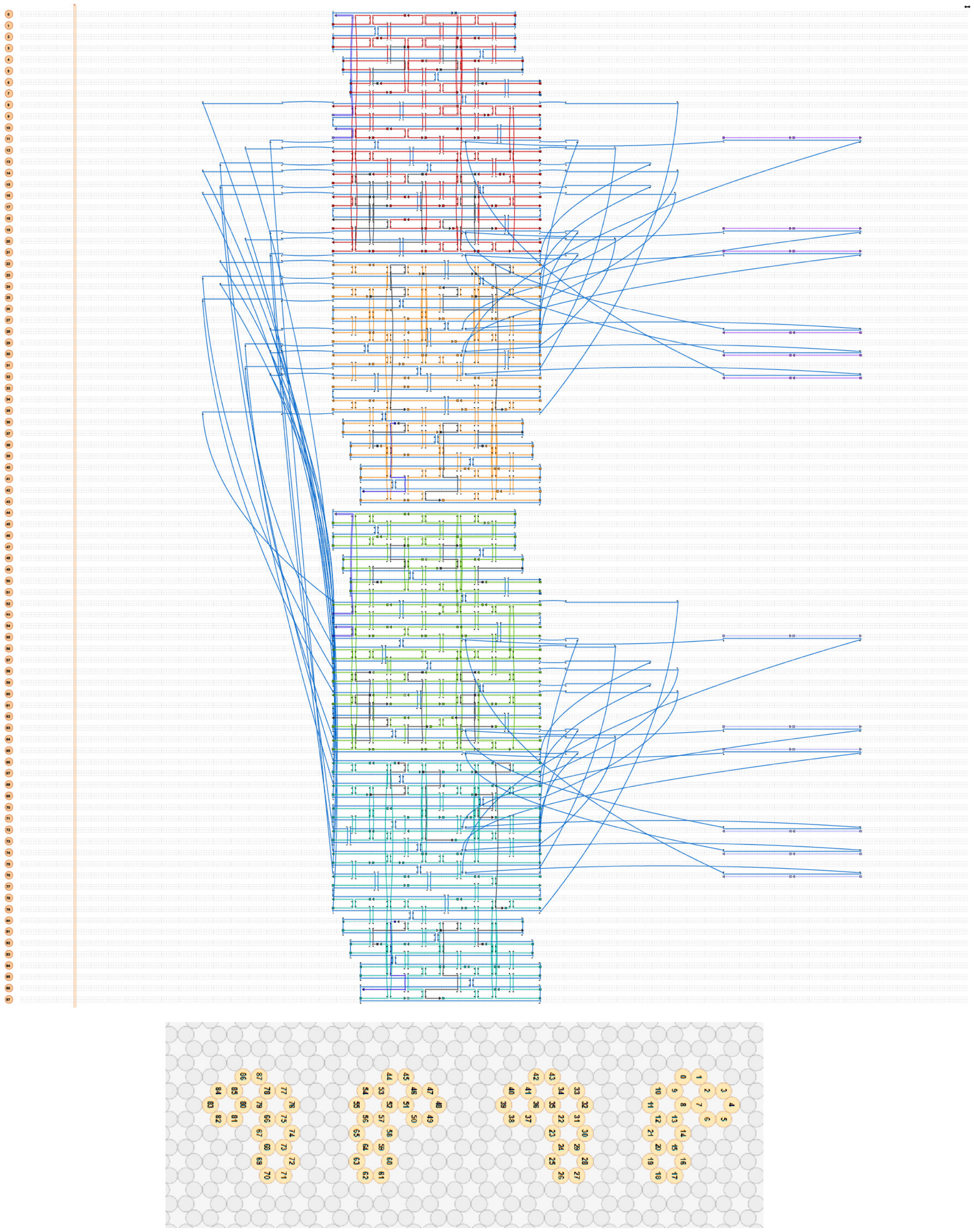

**Figure S1.** CaDNAno design file and depiction of the four 22-helix-bundle-subunits of the MechanoPore. The routing of the p8634 scaffold is shown in blue. Handles for cholesterol-modified oligonucleotides and handles for Cy5-functionalized oligonucleotides are depicted in black and dark blue, respectively. Additionally, the opening strands (in purple) are included in this image.

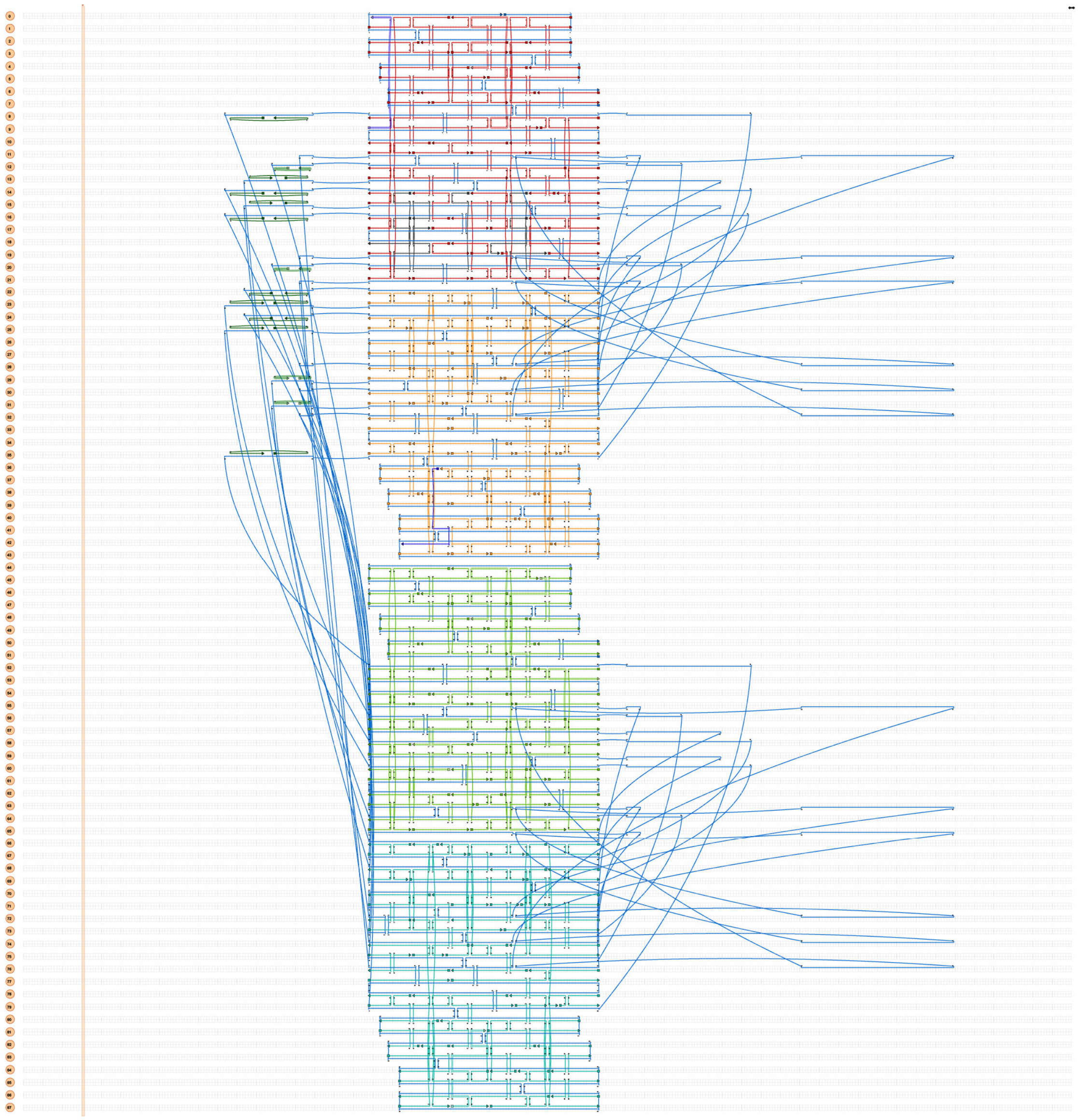

**Figure S2.** CaDNAno design file of the MechanoPore for DNA-PAINT measurements. Again, the routing of the p8634 scaffold is shown in blue. Handles for biotin-modified oligonucleotides and for PAINT imager strands are depicted in black and dark blue, respectively. Additionally, the closing strands (in dark green) are included in this image.

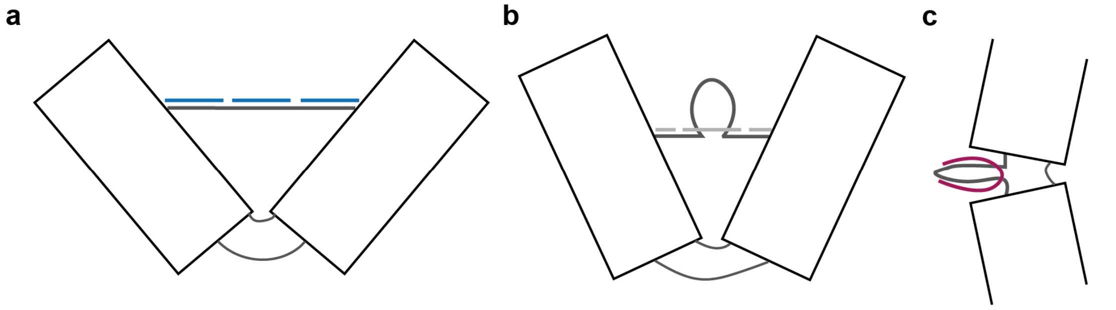


**Figure S3.** Schematic representation of the switching mechanism of the MechanoPore. **(a)** Fully open state: The opening of the MP is achieved via the addition of the opening strands (blue) that hybridize to the scaffold struts. These struts thus become double-stranded and 'push' the edges of the nanopores apart, leading to an opening of the nanopores. **(b)** Semi-open state: In order to achieve a different fixed opening angle of the MP, other trigger strands (grey) can be added whose binding results in a partially double-stranded strut with a single-stranded loop. By carefully choosing the length of the double-stranded regions, the opening angle of the nanopores can be tuned. **(c)** Closed state: For the closing mechanism, the closing strands (red) added to the sample that can bind to the linking scaffold parts in the corners without struts. The binding of these closing strands forces the neighbouring edges together, thereby leading to a decreased opening angle of the nanopores.

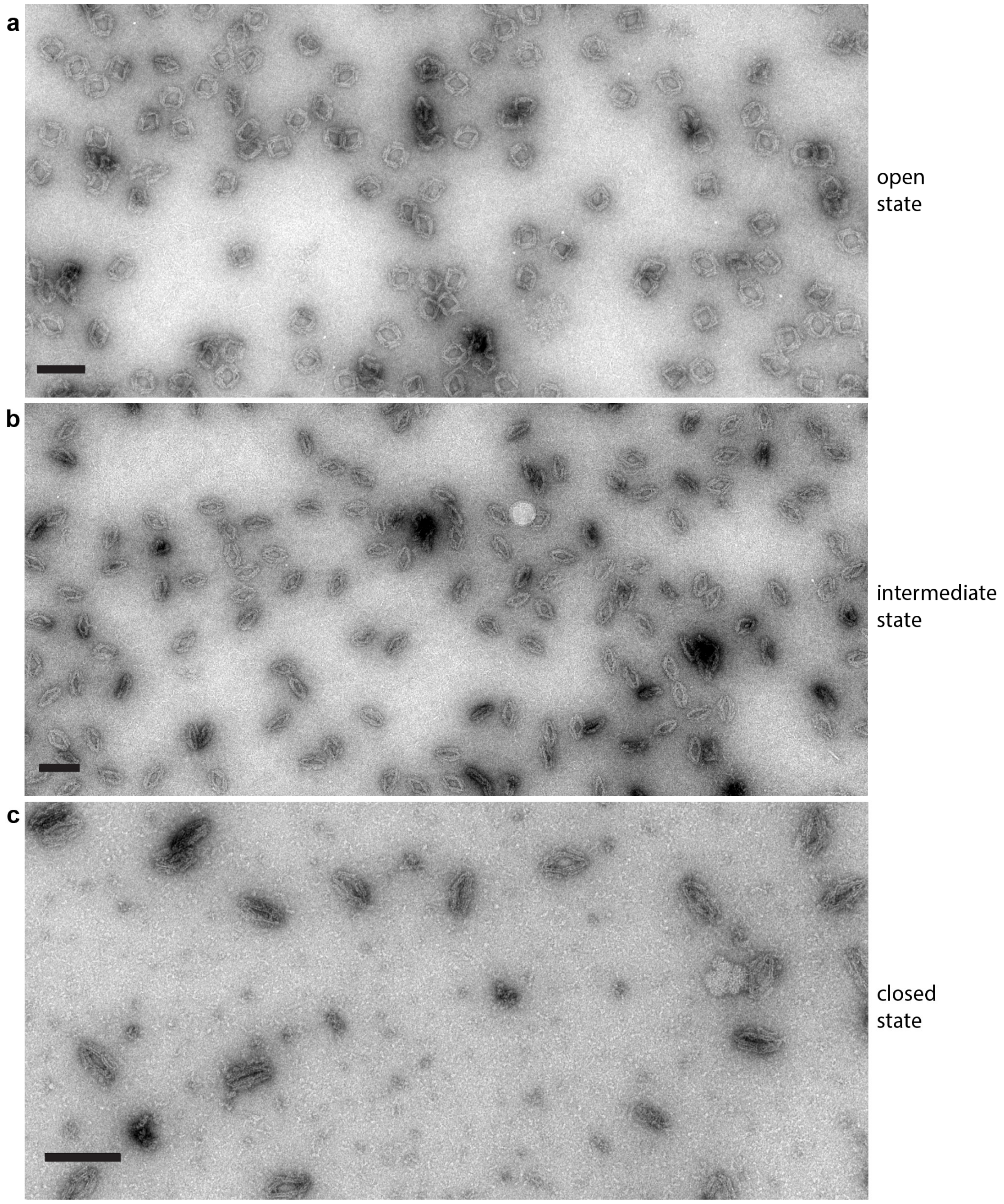


**Figure S4.** Low magnification TEM images of the MechanoPore in its three different configurations (a: open, b: intermediate and c: closed). The DNA origami nanopores are usually well-distributed and show only minor signs of aggregation or clustering. Mean opening angles were determined to be 89.1° ± 7.7° (open), 49.5° ± 7.5° (intermediate) and 25.6° ± 6.2° (closed). Scale bars: 100 nm.

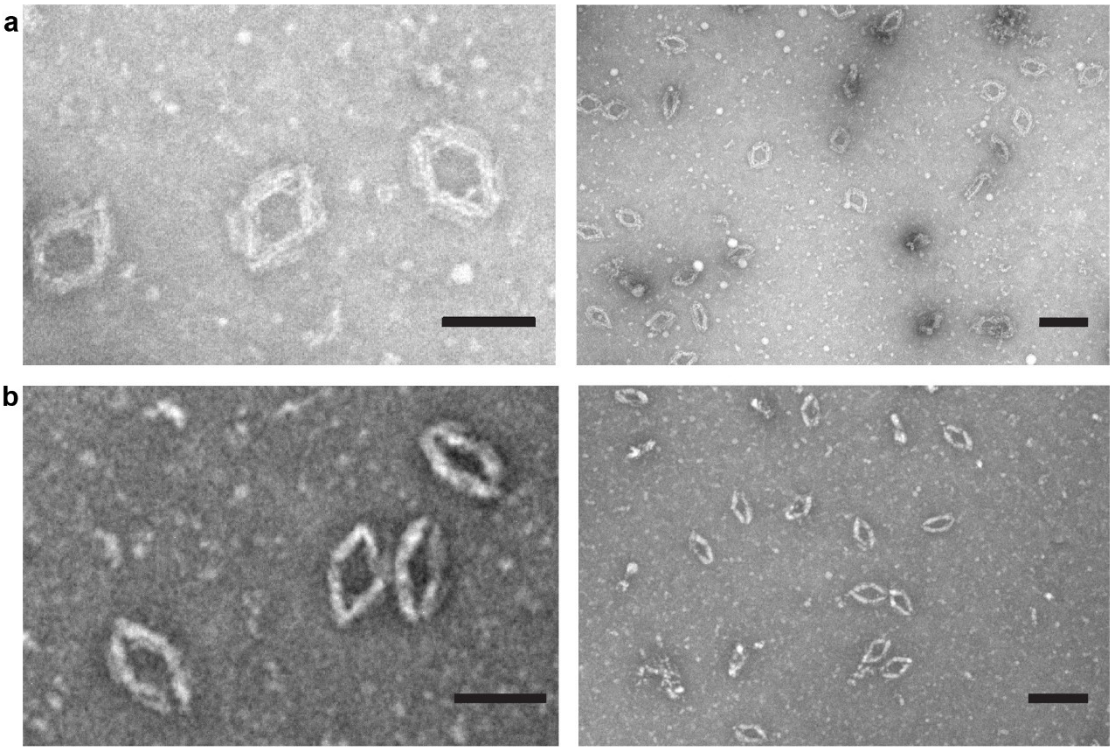


**Figure S5.** TEM images for two additional configurations that the MechanoPore can adopt via changing the nature of the opening strands. (a) 'Semi-open' state of the MechanoPore with an average opening angle of 68.7° ± 7.7°. (b) 'Fixed intermediate' state of the MechanoPore with an average opening angle of 52.3° ± 6.5°. Scale bars: 50 nm (left images) and 100 nm (right images).

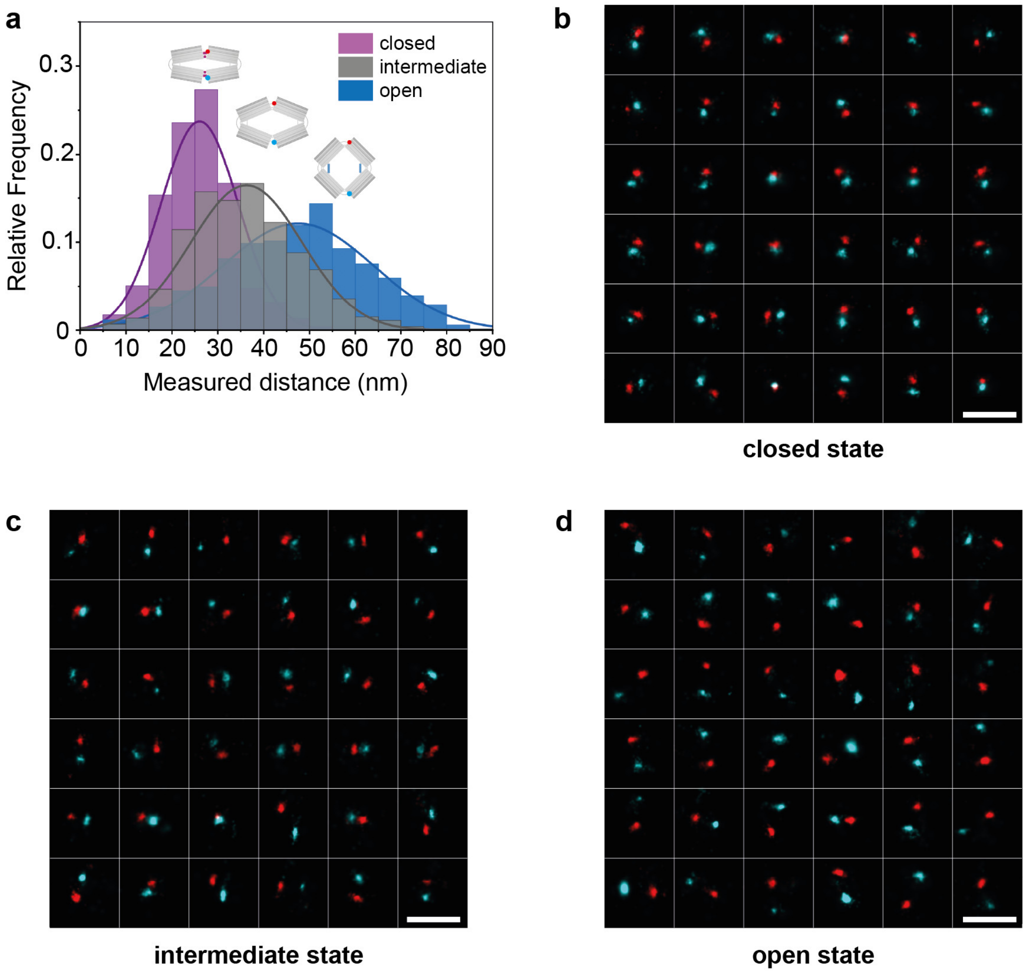


**Figure S6.** (a) Distance distributions obtained from 3D-DNA-PAINT measurements for MechanoPores in the three investigated configurations. For each state (closed, intermediate and open), around 700 - 750 nanopores were picked. A pronounced shift of the mean of the distributions for the different configurations is clearly visible. (b) – (d) Overview over 36 representative nanopores for the closed (b), intermediate (c) and open (d) state (red: R4-R3 imaging round, cyan: R6-R3 imaging round). Scale bars: 100 nm.

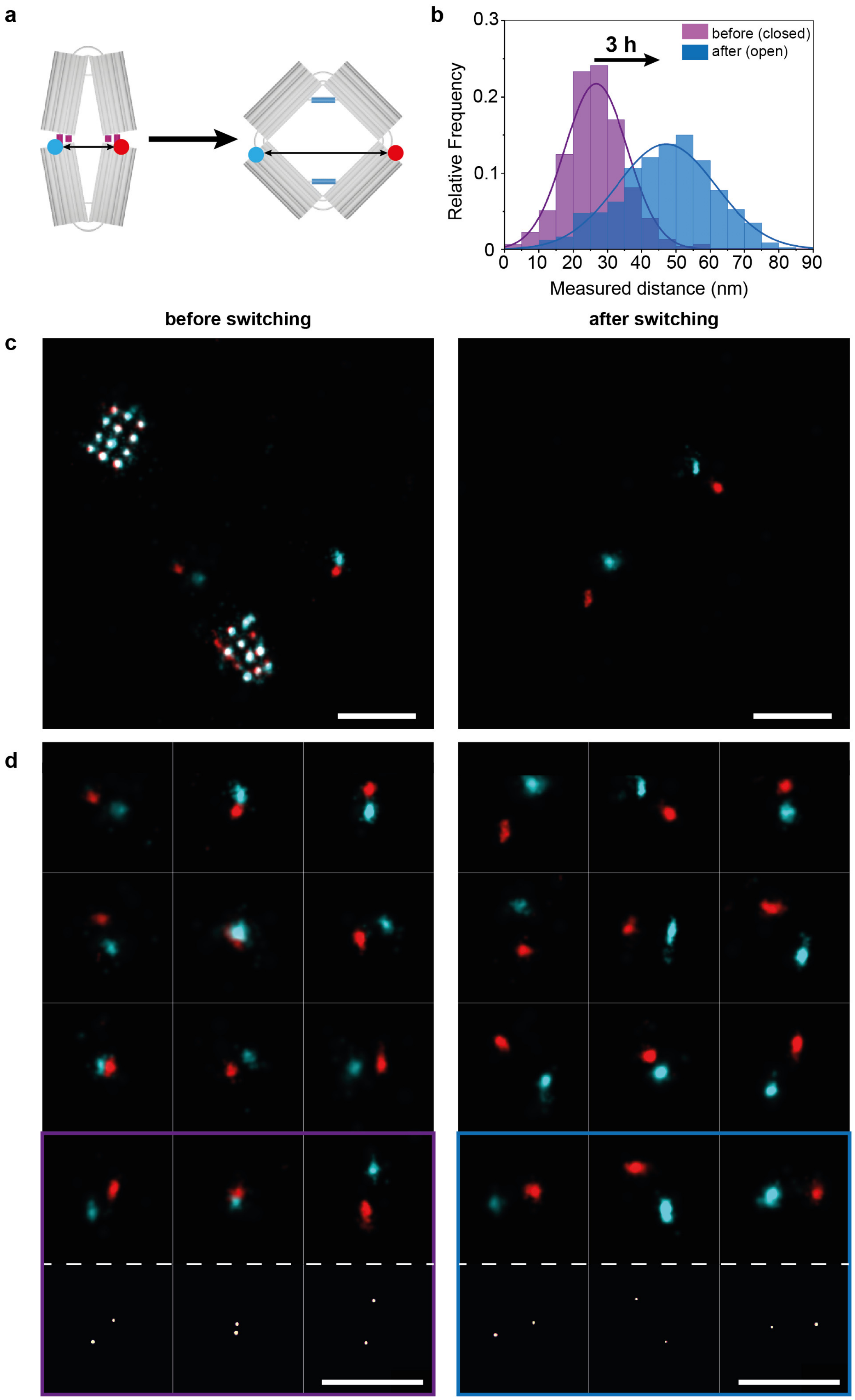


**Figure S7.** (a) Schematic representation of the configurational change of the MPs from the closed to the open state, investigated with the help of 3D-DNA-PAINT. (b) Distance distributions obtained from 3D-DNA-PAINT measurements before and after the switching of the nanopores induced by the addition of the corresponding trigger and anti-trigger strands (3 h incubation time). The pronounced shift of the mean of the distributions towards larger distances between the opposing corners clearly shows the successful configurational change of the nanopores. (c) Representative fields of view from the corresponding 3D-DNA-PAINT datasets. (d) Overview over representative selected nanopores. For distance measurements, localizations from each nanopore corner are grouped, yielding a more precise measure of the corners' coordinates and thus their distance (cluster centers in the bottom row). Scale bars: 100 nm.

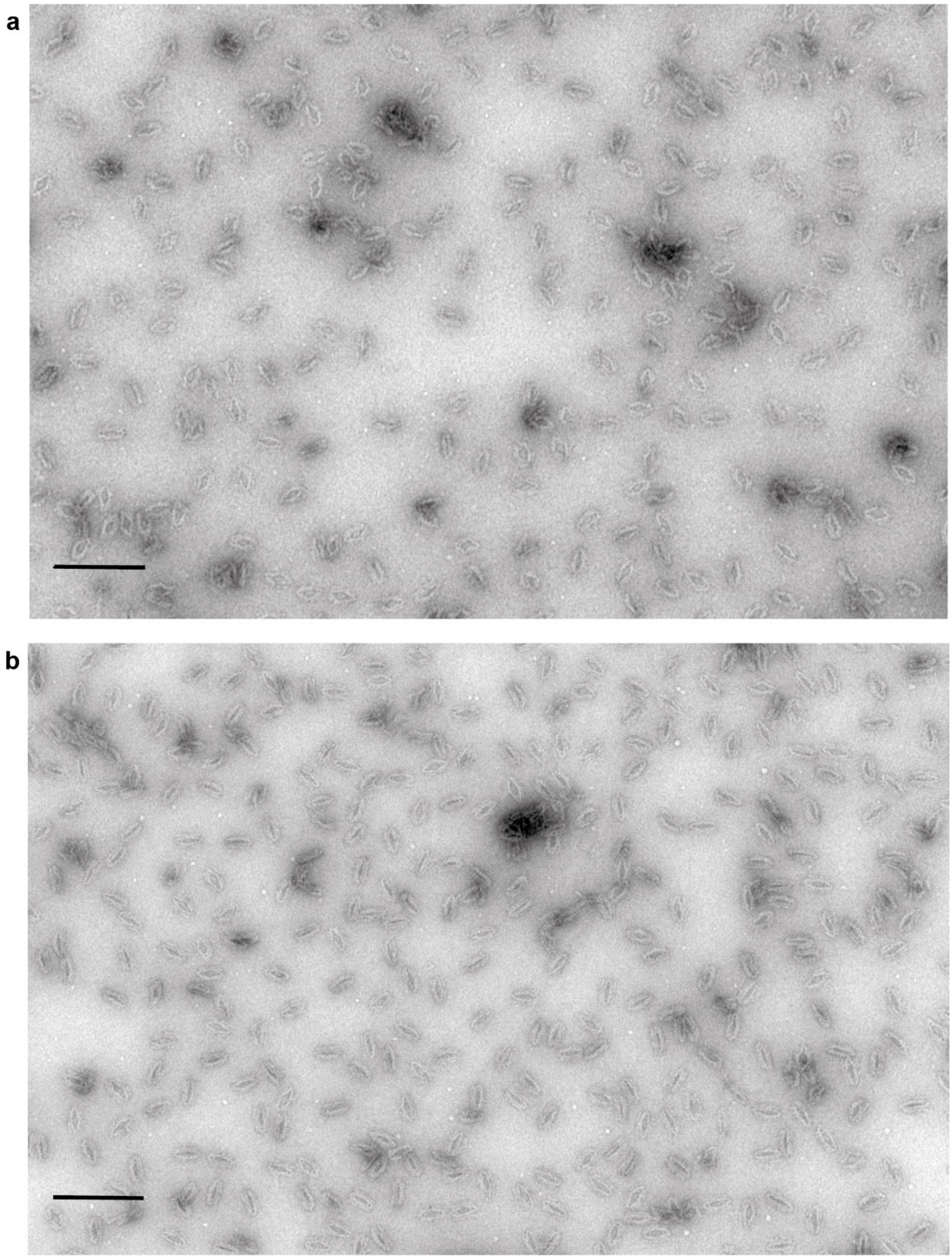

Figure S8. Low-magnification TEM images of MechanoPores in their (a) intermediate and (b) closed configurations with attached anti-handles with cholesterol-modifications. The images show that the addition of the cholesterol to the DNA origami nanostructures does not result in major aggregation or clustering and only few small aggregates are found in the samples. Scale bars: 200 nm.

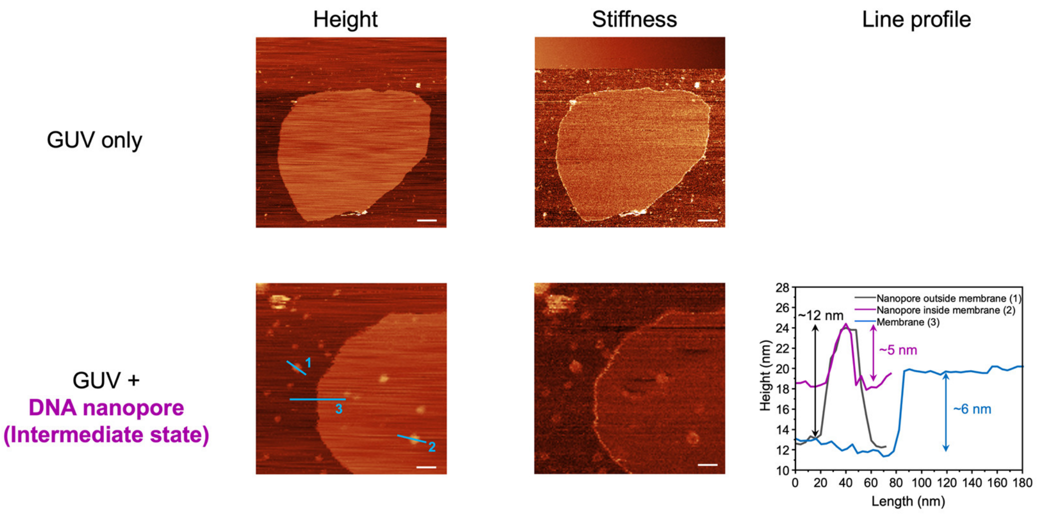


**Figure S9.** AFM images for blank GUV and GUV with MP (intermediate state) insertion. Here, the height and slope (stiffness) channels of QI$^{TM}$ (quantitative imaging) mode measurement are shown. The line profile plots the height of three lines on height channel of the GUV with DNA nanopore sample. Scale bars: 200 nm.

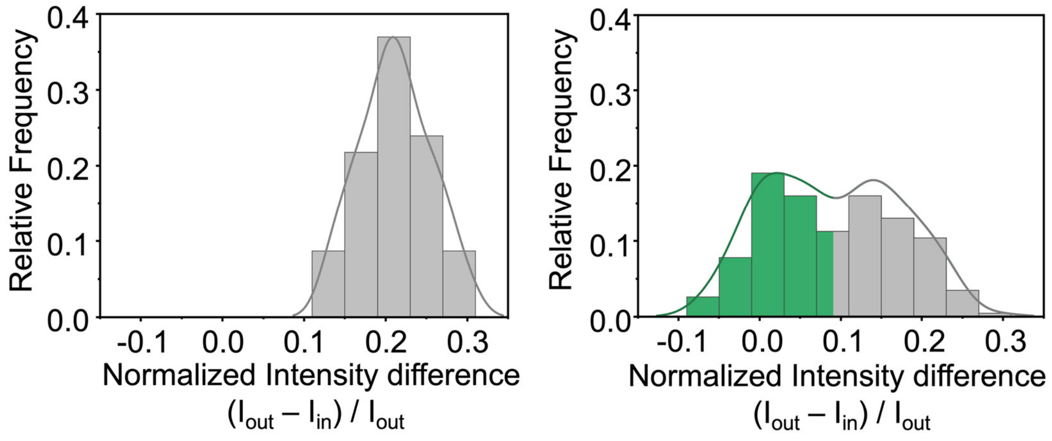


**Figure S10.** Statistical histograms and distribution curves of the normalized intensity difference $I_{ndiff} = (I_{out} - I_{in}) / I_{out}$ for MP-free GUVs (left, N = 69) and GUVs containing MP-I (right, N = 118), respectively. $I_{out}$ and $I_{in}$: fluorescence intensities outside and inside the GUV boundary, respectively. Green: filled vesicles ($I_{ndiff} < 0.09$). Grey: unfilled vesicles ($I_{ndiff} \geq 0.09$).

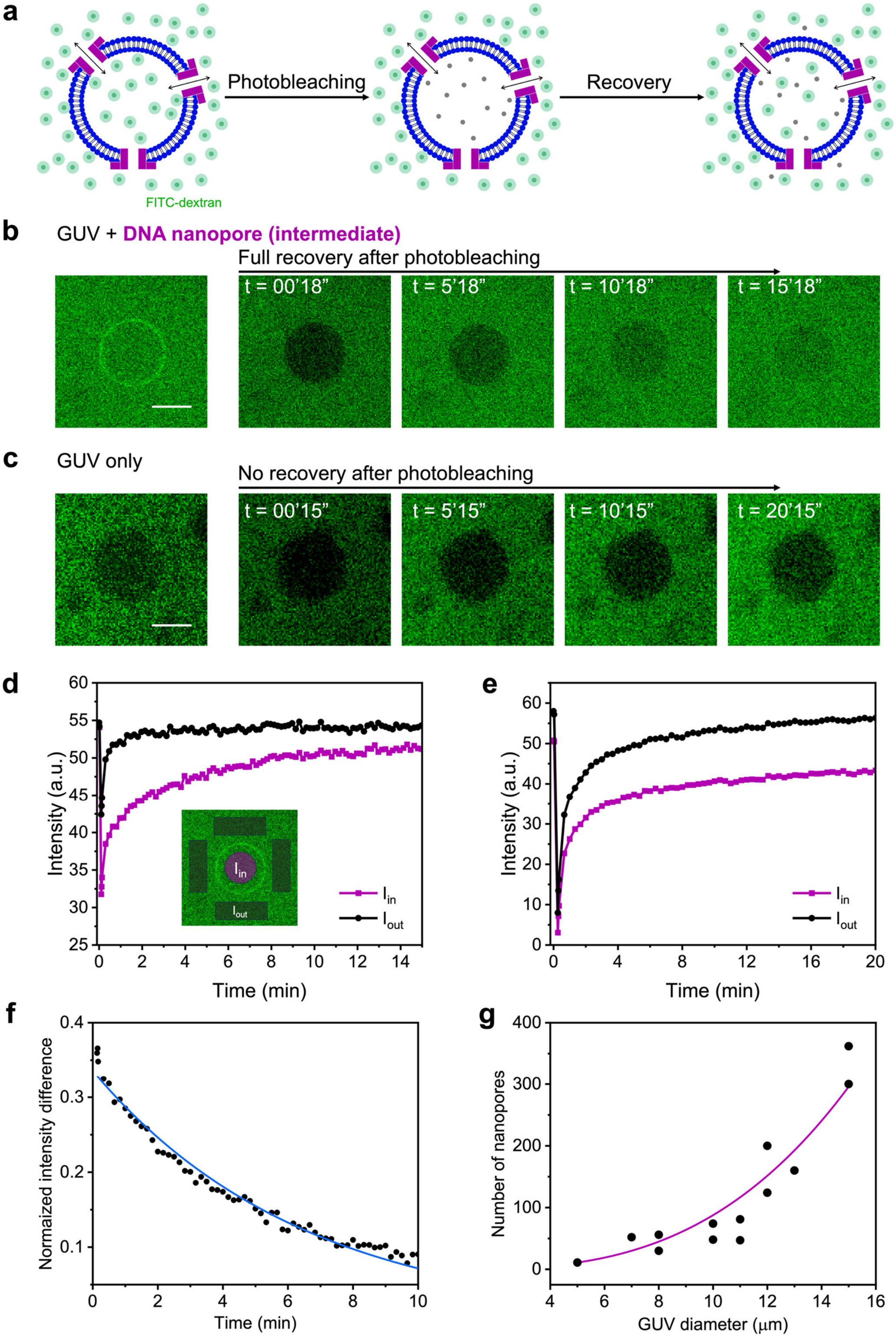


**Figure S11.** Diffusion of dextran across MPs. (a) Schematic representation of the FRAP assay on the GUV with MP in the intermediate state. (b) and (c) Representative FRAP images of GUV only and GUV with MP in the intermediate state in the presence of 70 kDa FITC-dextran. (d) and (e) FRAP traces of GUV only and GUV with MP in the intermediate state in the presence of 70 kDa FITC-dextran. Scale bars: 10 µm. Inset: Indication of fluorescence intensity inside ($I_{in}$) and outside ($I_{out}$) GUVs. (f) Normalized intensity difference $I_{ndiff}$ over time for the GUV illustrated in (b) and (d). Data are fitted by diffusion model. For such a 12 µm diameter vesicle, we estimated a number of functional pores $N_p$ = 125. (g) Number of DNA nanopores extracted from fits for different GUVs, as a function of GUV diameter (black circle). Averaged data show an increase of the number of pores as a function of GUV diameter. Magenta line represents an allometric fitting (3rd order) to the data.

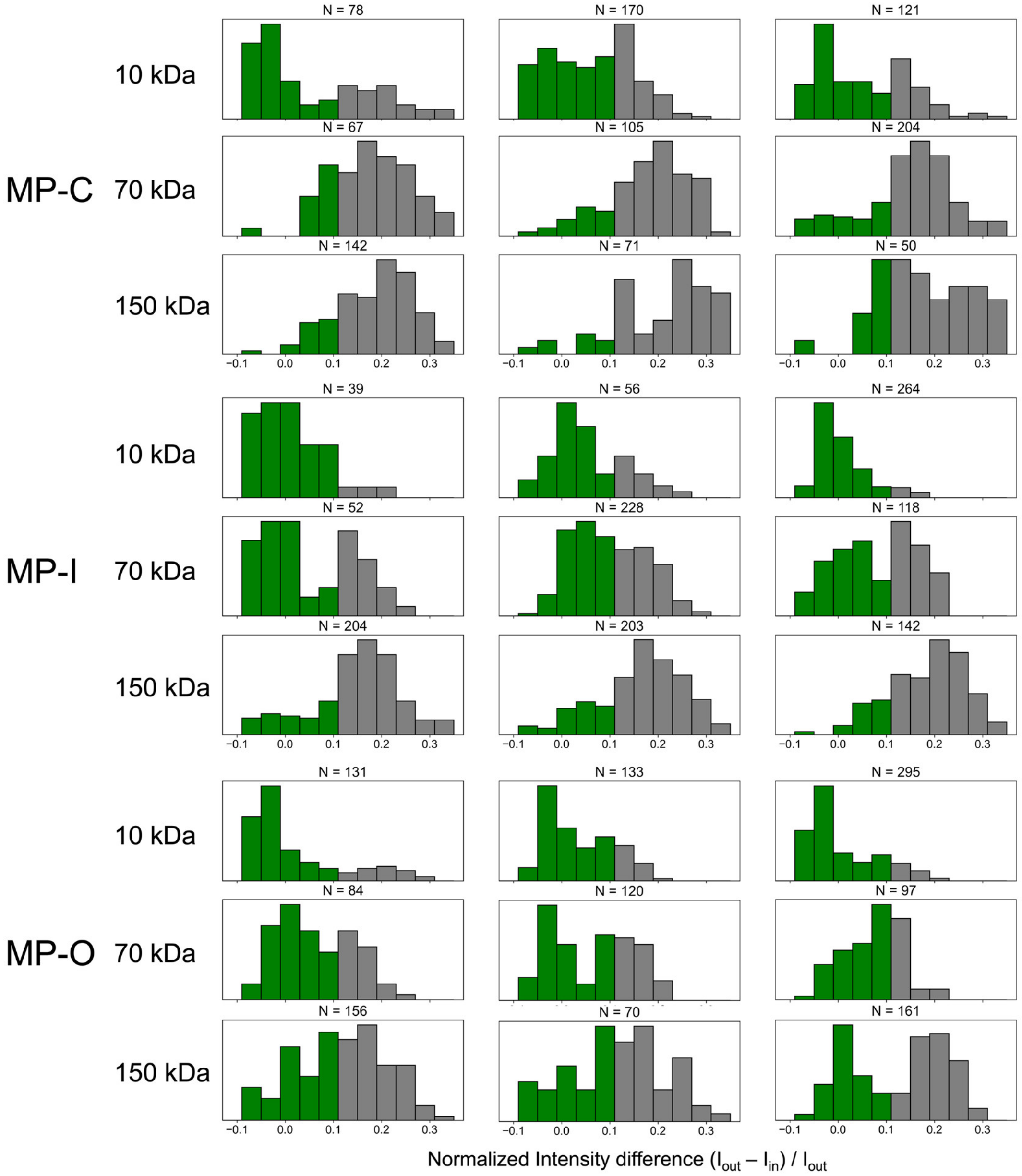


**Figure S12.** Statistical histograms of the normalized intensity difference $I_{ndiff} = (I_{out} - I_{in}) / I_{out}$ for GUVs containing MP-C, MP-I, or MP-O with 10 kDa, 70 kDa, or 150 kDa FITC-dextran. $I_{out}$ and $I_{in}$: fluorescence intensities outside and inside the GUV boundary, respectively. The N numbers above each subfigure represent the numbers of GUVs measured.

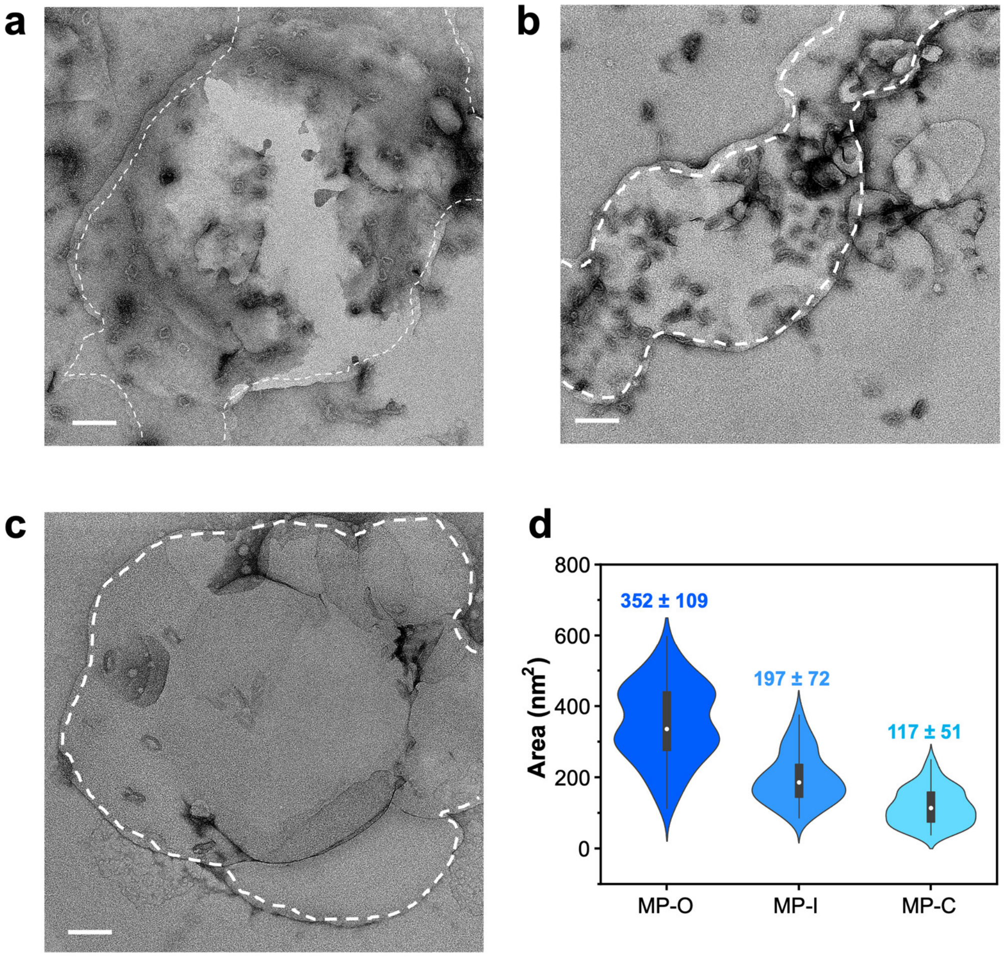


**Figure S13.** TEM images for the MechanoPores in three states (a: open, b: intermediate, and c: closed) after embedding into a lipid membrane in a large view. (d) Statistical analyses of the inner area of the DNA origami nanopores in different states after membrane insertion. Scale bars: 100 nm.

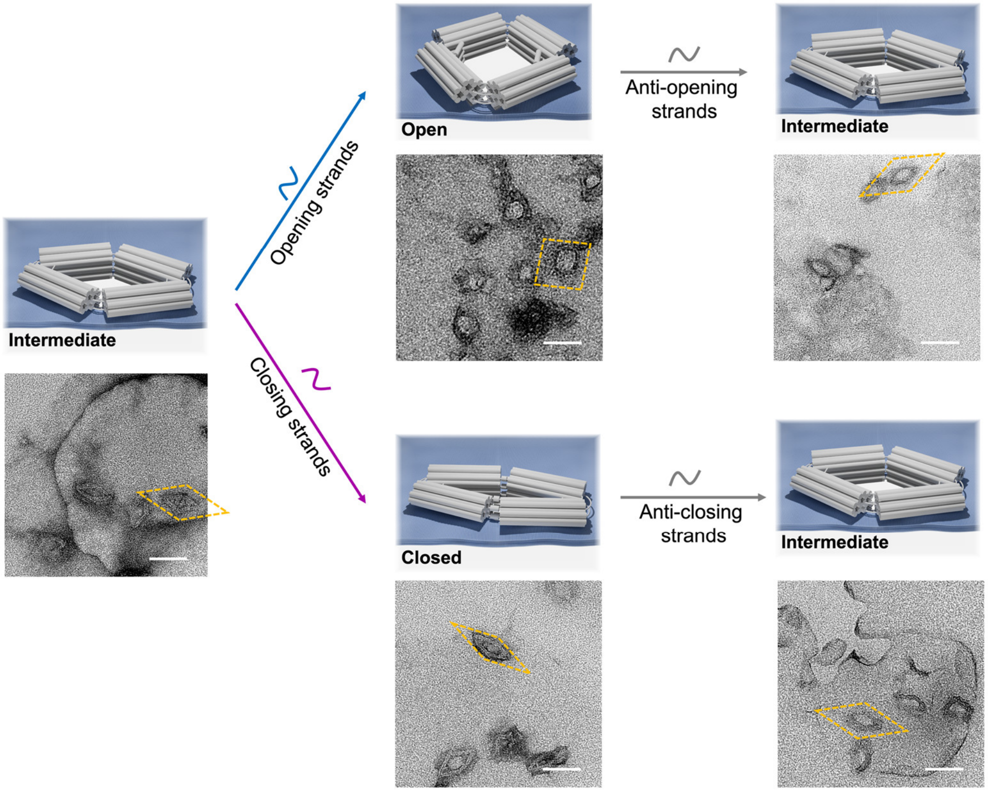


**Figure S14.** TEM images of conformational changes of MechanoPores after membrane insertion. Schematic representation of the process of reversible conformational changes of the DNA nanoactuator embedded in a lipid bilayer membrane. The procedure starts from the intermediate state, followed by the addition of the opening or closing strands. Subsequently, the corresponding anti-trigger/releasing strands are added. TEM images show the corresponding states of the nanopores after switching. Scale bars: 50 nm.

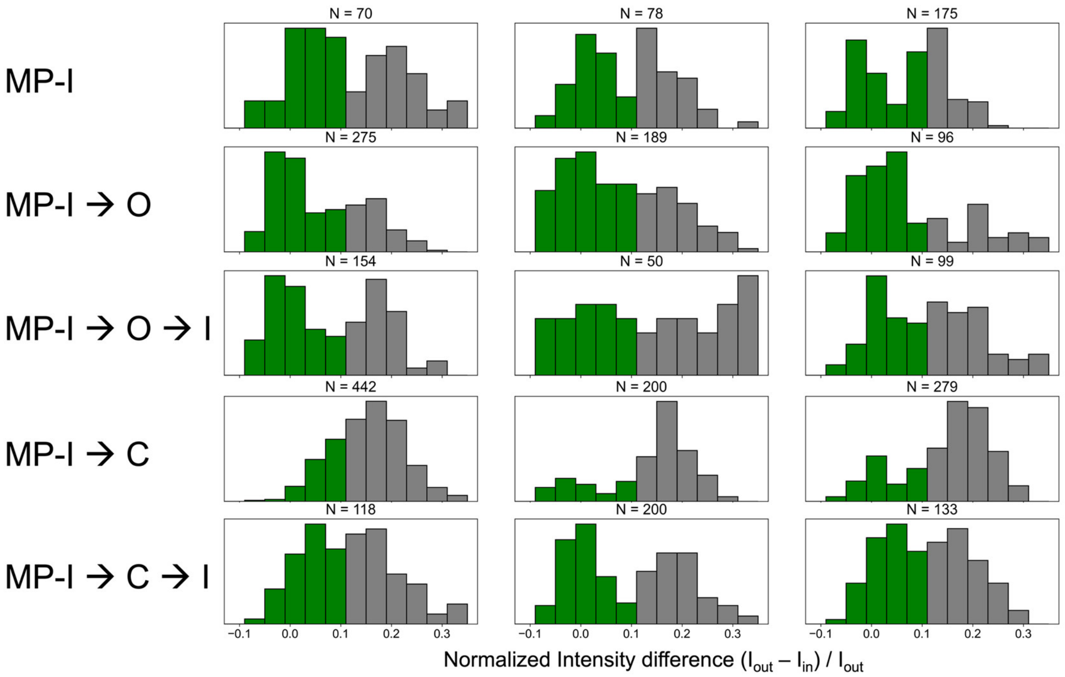


**Figure S15.** Statistical histograms of the normalized intensity difference $I_{ndiff} = (I_{out} - I_{in}) / I_{out}$ for reversible conformational changes of MPs embedded in GUVs with 70 kDa FITC-dextran. $I_{out}$ and $I_{in}$: fluorescence intensities outside and inside the GUV boundary, respectively. The N numbers above each subfigure represent the numbers of GUVs measured.

Table S1: Sequences of the handles and modified anti-handles that were incorporated into the MechanoPores and of the trigger strands employed for the opening or closing of the nanopores. Anti-trigger strands are obtained by using the reverse complement of the trigger strand sequences. Binding sites for the anti-handles with Cy5 or cholesterol/biotin modifications are marked in blue or green, respectively. The toeholds of the opening and closing strands that are used for the displacement of these staples are marked in red.

| **Name** | **Sequence** |
|---|---|
| Handle for Cy5 attachment 1 | AATTATTCATTCACCAGCTTGAGGACTTAAAA |
| Handle for Cy5 attachment 2 | AGACACATAGTCGGTGCAAAATCACCAGCTTGAGGACTTAAAA |
| Handle for Cy5 attachment 3 | CCGAGGAAAGCAAGCCGCTTGAGGACTTAAAA |
| Handle for Cy5 attachment 4 | CCCGTATAAAGAAGGAACTTGAGGACTTAAAA |
| Handle for Cy5 attachment 5 | AGCCTTGCTGGTTGAAAGGAATTGAGGACTTGAGGACTTAAAA |
| Handle for Cy5 attachment 6 | CACACGACCGGTGCCGTCTTGAGGACTTAAAA |
| Cy5-modified anti-handle | Cy5 - TTTTAAGTCCTCAAG |
| DNA-PAINT handle 1 | CACACGACCGGTGCCGTAACAACAACAACAACAACAA |
| DNA-PAINT handle 2 | AGACACATAGTCGGTGCAAAATCACCAG ACACACACACACACACACA |
| DNA-PAINT docking site 20 nm grid | CTCTCTCTCTCTCTCTC |
| DNA-PAINT imager 5xR6 | TGTTGTT – Cy3B |
| DNA-PAINT imager 7xR4 | GTGTGT – Cy3B |
| DNA-PAINT imager 7xR3 | GAGAGAG – Cy3B |
| Handle for cholesterol attachment 1 | TATGAGAAGTTAGGAATGTTATTTTGATATATAGTTTCCG CTTTTGCCAATACTGCGGAAGTAATCTACGTAAC |
| Handle for cholesterol attachment 2 | TATGAGAAGTTAGGAATGTTATTTTAACCATCATACGTATT CAGGGATAAATAGCGAGAGATTAAACGCTTGCC |
| Handle for cholesterol | TATGAGAAGTTAGGAATGTTATTTTTCTACTAGCACCAACT |

| | |
|---|---|
| attachment 3 | CATGCCATTTGAAAGAGGA |
| Handle for cholesterol attachment 4 | TATGAGAAGTTAGGAATGTTATTTTCTGACGACAAATCATGACAAGAACCGGATA |
| Handle for cholesterol attachment 5 | TATGAGAAGTTAGGAATGTTATTTTAAACGAATTGACCCGACCAGGCGCATAGTCCC |
| Handle for cholesterol attachment 6 | TATGAGAAGTTAGGAATGTTATTTTGCCCGGAATACGGGAGAAGCC |
| Handle for cholesterol attachment 7 | TATGAGAAGTTAGGAATGTTATTTTTTTAGATTTAGTTTGAACATTACAGT |
| Handle for cholesterol attachment 8 | TATGAGAAGTTAGGAATGTTATTTTGCGCCATTCGCCAGACCCTCATTGATATACAGGAGGAAAGCGC |
| Handle for cholesterol attachment 9 | TATGAGAAGTTAGGAATGTTATTTTTCAGGATTGACTTCAAATATCGCGTTTTAAAACTCCCAAGGAT |
| Handle for cholesterol attachment 10 | TATGAGAAGTTAGGAATGTTATTTTAGCCGCCGAAACAAATTTGCCAAGCGCGTTTTCATCAAACGTC |
| Handle for cholesterol attachment 11 | TATGAGAAGTTAGGAATGTTATTTTCCATAAGTTTATTTTGTAGCC |
| Handle for cholesterol attachment 12 | TATGAGAAGTTAGGAATGTTATTTTGCCTGACAGGAGGTTGAGGCAG |
| Handle for cholesterol attachment 13 | TATGAGAAGTTAGGAATGTTATTTTAGTAATAACGACTTGACGCGAG |
| Handle for cholesterol attachment 14 | TATGAGAAGTTAGGAATGTTATTTTTGAGAGAGCTAGGGCGGGAAGA |
| Handle for cholesterol attachment 15 | TATGAGAAGTTAGGAATGTTATTTTAACACAATTTTATCCTGGCCTTAAGGCTTAT |
| Handle for cholesterol attachment 16 | TATGAGAAGTTAGGAATGTTATTTTCAATAAAGTACCGACAAAAGGT |
| Handle for cholesterol attachment 17 | TATGAGAAGTTAGGAATGTTATTTTACTCCTCGCCGGAGGAAGTGAAACTTATCAGTAAAGGGATAGGTGCATC |
| Handle for cholesterol attachment 18 | TATGAGAAGTTAGGAATGTTATTTTTGAAAGTATCAATAACGAAGTGGAAACCAG |
| Handle for cholesterol attachment 19 | TATGAGAAGTTAGGAATGTTATTTTGGTATAGTCTAGCTGTATTGCCGTAAATGTGAGCG |
| Handle for cholesterol attachment 20 | TATGAGAAGTTAGGAATGTTATTTTAACAATGCGCCGCTACACGGTCACGCGAACG |
| Handle for cholesterol attachment 21 | TATGAGAAGTTAGGAATGTTATTTTTGCAAGCGGTCCACGCTGGTTT |
| Handle for cholesterol | TATGAGAAGTTAGGAATGTTATTTTCCTCAGGCGTAACC |

| | |
|---|---|
| attachment 22 | GTCACGTTGGTGTAGA |
| Handle for cholesterol attachment 23 | TATGAGAAGTTAGGAATGTTATTTTAATTAATCAAAGG CTTCTCCGTG |
| Handle for cholesterol attachment 24 | TATGAGAAGTTAGGAATGTTATTTTAGTTTATAGCTGA AAGTAGGGAAACTG |
| Handle for cholesterol attachment 25 | TATGAGAAGTTAGGAATGTTATTTTGAGTTAGAGTCTG AGTATCACGATAA |
| Handle for cholesterol attachment 26 | TATGAGAAGTTAGGAATGTTATTTTATTGTGAATTACCTT CAGCATCAGACATTAAAC |
| Handle for cholesterol attachment 27 | TATGAGAAGTTAGGAATGTTATTTTACCTGTCTTATGA AGCGGGGTT |
| Handle for cholesterol attachment 28 | TATGAGAAGTTAGGAATGTTATTTTGAAGGGTTCGTAGAT AACCCTCTGGTCAGTTGGCAACTAACAA |
| Handle for cholesterol attachment 29 | TATGAGAAGTTAGGAATGTTATTTTGCCAACCACCAGC AGAATGAT |
| Handle for cholesterol attachment 30 | TATGAGAAGTTAGGAATGTTATTTTGATACCATATCAA AATTATTTG |
| Cholesterol-modified anti-handle | TAACATTCCTAACTTCTCATA – chol-TEG |
| Handle for biotin attachment 1 | TATGAGAAGTTAGGAATGTTATTTTGATATATAGTTTCCGC TTTTGCCAATACTGCGGAAGTAATCTACGTAAC |
| Handle for biotin attachment 2 | TATGAGAAGTTAGGAATGTTATTTTAACCATCATACGTATT CAGGGATAAATAGCGAGAGATTAAACGCTTGCC |
| Handle for biotin attachment 3 | TATGAGAAGTTAGGAATGTTATTTTTCTACTAGCACCAACT CATGCCATTTGAAAGAGGA |
| Handle for biotin attachment 4 | TATGAGAAGTTAGGAATGTTATTTTCTGACGACAAATCAT GACAAGAACCGGATA |
| Handle for biotin attachment 5 | TATGAGAAGTTAGGAATGTTATTTTAAACGAATTGACCCG ACCAGGCGCATAGTCCC |
| Biotin-modified anti-handle | TAACATTCCTAACTTCTCATA – biotin-TEG |

| **Name** | **Sequence** |
|---|---|
| Opening strand 1 | ATCCCCTCCGGGTACCGAGCTCGAATTCGTAATCA |
| Opening strand 2 | TGGTCATAGCTGTTTCCTGTGTGAAATTTCTCTAA |

| | |
|---|---|
| Opening strand 3 | CCACATTAATGCTTTAAACAGTTCAGAAAACGAGA |
| Opening strand 4 | ATGACCATAAATCAAAAATCAGGTCTTTAATCCAA |
| Opening strand 5 | TTCTCTCATTCAGCATCAGAAATAGAAGAATTACA |
| Opening strand 6 | GCGCAACACAGCAATAAAAATGCGCCGCATTTACC |
| Opening strand 7 | CAATCATAAGGGAACCGAACTGACCAACACCCTAC |
| Opening strand 8 | TAACTCATACTTAGCCGGAACGAGGCGCAGACGGT |
| Opening strand 9 | GGAGTGAGAATAGAAAGGAACAACTAAATCATTCC |
| Opening strand 10 | TCTTTAATTGCTAAACAACTTTCAACAGTTTCAGC |
| Opening strand 11 | GAACAATATTACCGCCAGCCATTGCAACATCCCAC |
| Opening strand 12 | CATTCCTAAACTATCGGCCTTGCTGGTAATATCCA |
| Opening strand 13 | CCACTAACCTCCGGCTTAGGTTGGGTTATATAACT |
| Opening strand 14 | ATATGTAAATGCTGATGCAAATCCAATCACTTCAT |
| Opening strand 15 | ACTCCTACCATCCTGGGAAGACTCCTGTTATCAAG |
| Opening strand 16 | CACTGCACTGGTGACCTGGAAGAGTTTCCCCATAC |
| Opening strand 17 | CACCTTTACTGCTGGATGAACGGGAAAGAAACCAG |
| Opening strand 18 | CAATACATCAAACGCCGCGACCAGGAGAACCTACT |
| Opening strand 19 | CCTTCCTGTAGCCAGCTTTCATCAACATTTAATCA |
| Opening strand 20 | CAACTCCAACGCCATCAAAAATAATTCGCGTCTGG |
| Opening strand 21 | AGATTGTATAAGCAAATATTTAAATTGTTACCCAC |
| Opening strand 22 | CCAATATGATAATCAGAAAAGCCCCAAAAACAGGA |
| Opening strand 23 | GTTAAGCCCAATAATAAGAGCAAGAAACCCTTACC |
| Opening strand 24 | TATAACATATCAGAGAGATAACCCACAAGAATTGA |

| | |
|---|---|
| Closing strand 1 | CTCACCCCTCGCGGGGCCCAACAATCCT |
| Closing strand 2 | CTATCCTAAGGCCGCTTTTGTTTGGCTTGCAGGGAG |
| Closing strand 3 | TTCATACTAAAGACTTCCCAGGCTTTGA |
| Closing strand 4 | CCTTATCGCTTGAGATGGTTCCCACGAGTAGTAAAT |
| Closing strand 5 | CCACCATGTGGCACAGACAAGGGGAAAGCGTAAGAA |
| Closing strand 6 | AAACTCACGCCACCCTTTTCCACCCTCA |
| Closing strand 7 | CAACTATTTGCTCAGTACCACCCTAGGATTAGCGGG |
| Closing strand 8 | ACTACATGATTCAAAACCCATGTGTAGG |
| Closing strand 9 | TCAACAACACCGCTTCTGGTAAACCAGCCAGCTTTC |
| Closing strand 10 | TACCCTTGCAAACGTAGAAACCCATTACGCAGTATG |
| Closing strand 11 | AATCTACACATCATTTTGAAA |
| Closing strand 12 | ATCTTACAGGGGGAAATTTGC |
| Closing strand 13 | ACAATTTACTGCTTTCTCTCA |
| Closing strand 14 | CTTATAACGTAATTTCCCATG |
| Opening strand 1 for ‘semi-open’ state | ATCCCCTCCGGGTACCGAGCTCGAATTGCTGTTTCCTGTGTGAAATT |
| Opening strand 2 for ‘semi-open’ state | TCTCTAAAATGCTTTAAACAGTTCAGAAAATCAAAAATCAGGTCTTT |
| Opening strand 3 for ‘semi-open’ state | CCACATTATTCAGCATCAGAAATAGAACAGCAATAAAAATGCGCCGC |
| Opening strand 4 for ‘semi-open’ state | AATCCAATACTTAGCCGGAACGAGGCGAGGGAACCGAACTGACCAAC |

| | |
|---|---|
| Opening strand 5 for 'semi-open' state | TTCTCTCTTGCTAAACAACTTTCAACAAATAGAAA GGAACAACTAAA |
| Opening strand 6 for 'semi-open' state | ATTTACCAAACTATCGGCCTTGCTGGTTTACCGCCAG CCATTGCAAC |
| Opening strand 7 for 'semi-open' state | ACCCTACCCTCCGGCTTAGGTTGGGTTATGCTGATG CAAATCCAATC |
| Opening strand 8 for 'semi-open' state | TAACTCACCATCCTGGGAAGACTCCTGTGGTGACCT GGAAGAGTTTC |
| Opening strand 9 for 'semi-open' state | TCATTCCACTGCTGGATGAACGGGAAACAAACGCC GCGACCAGGAGA |
| Opening strand 10 for 'semi-open' state | TCTTTAAAACGCCATCAAAAATAATTCTAGCCAGCTT TCATCAACAT |
| Opening strand 11 for 'semi-open' state | ATCCCACGATAATCAGAAAAGCCCCAATAAGCAAAT ATTTAAATTGT |
| Opening strand 12 for 'semi-open' state | CATTCCTTATCAGAGAGATAACCCACACAATAATAAG AGCAAGAAAC |
| Opening strand 1 for 'fixed intermediate' state | ATCCCCTCCGGGTACCGAGCTTCCTGTGTGAAATT |
| Opening strand 2 for 'fixed intermediate' state | TCTCTAAAATGCTTTAAACAGAAAATCAGGTCTTT |

| | |
|---|---|
| Opening strand 3 for 'fixed intermediate' state | CCACATTATTCAGCATCAGAATAAAAATGCGCCGC |
| Opening strand 4 for 'fixed intermediate' state | AATCCAATACTTAGCCGGAACCCGAACTGACCAAC |
| Opening strand 5 for 'fixed intermediate' state | TTCTCTCTTGCTAAACAACTTAAGGAACAACTAAA |
| Opening strand 6 for 'fixed intermediate' state | ATTTACCAAACTATCGGCCTTCCAGCCATTGCAAC |
| Opening strand 7 for 'fixed intermediate' state | ACCCTACCCTCCGGCTTAGGTATGCAAATCCAATC |
| Opening strand 8 for 'fixed intermediate' state | TAACTCACCATCCTGGGAAGACCTGGAAGAGTTTC |
| Opening strand 9 for 'fixed intermediate' state | TCATTCCACTGCTGGATGAACCCGCGACCAGGAGA |
| Opening strand 10 for 'fixed intermediate' state | TCTTTAAAACGCCATCAAAAAGCTTTCATCAACAT |
| Opening strand 11 for 'fixed | ATCCCACGATAATCAGAAAAGAATATTTAAATTGT |

| | |
|---|---|
| intermediate' state | |
| Opening strand 12 for 'fixed intermediate' state | CATTCCTTATCAGAGAGATAATAAGAGCAAGAAAC |
